# Single-cell map of the female brain across reproductive transitions

**DOI:** 10.64898/2026.01.03.697507

**Authors:** Luisa Demarchi, Maria Tickerhoof, Ayyappa Kumar Sista Kameshwar, Devin Rocks, Laila Ouldibbat, Ana Milosevic, Masako Suzuki, Marija Kundakovic

## Abstract

Ovarian hormone shifts enable reproduction and are associated with substantial brain plasticity and disease risks. While imaging studies provide (micro)structural insights into brain changes across the ovarian cycle and pregnancy, the high resolution, single-cell map of the brain across reproductive transitions is missing. Here, we performed multiome (gene expression and chromatin accessibility) analysis of the mouse ventral hippocampus (vHIP) across sex, estrous cycle, and peripartum period at single cell resolution. We identify dynamic changes in vHIP cellular composition across the estrous cycle and pregnancy, including in the neural stem cells of the dentate gyrus (DG), enabling hormone-driven neurogenesis. Major gene expression changes are neuronal function-relevant, cell type-specific, and found in excitatory neurons of CA1, CA3, and DG subfields, across sex and reproductive transitions. In contrast, chromatin accessibility changes are more extensive and found across cell types, likely driven by estrogen level shifts in both within-female and between-sex comparisons. We show that chromatin remodeling during the estrous cycle primes the genome for gene expression changes during pregnancy and is also enriched for brain disease-relevant genes. Finally, we reveal a thyroid hormone transporter (Transthyretin, *Ttr*) gene as the major candidate gene that drives structural and behavioral changes across the estrous cycle and pregnancy. Our study provides an extensive cellular and molecular view of how reproductive transitions shape the brain and opens the possibility to target downstream targets of estrogen, including thyroid hormone signaling, as a treatment option for hormone-sensitive periods in women.

---

Ovarian hormone fluctuations are required for female reproductive function and occur systemically across the ovarian cycle as well as during pregnancy and postpartum. While hormone-driven remodeling of reproductive organs plays an important role in preparation for pregnancy and giving birth, the brain also undergoes significant changes with hormone shifts, representing a particular form of sex-specific brain plasticity that has been understudied, despite its critical importance for understanding female brain physiology and disease risk.

Studies in the early 90s showed that hormone fluctuations across the estrous cycle induce changes in dendritic spine density in the rodent hippocampus^1^, primarily driven by estrogen level changes^2^. Recently, high resolution imaging studies in humans showed gray matter changes across the menstrual cycle^3^ and in pregnancy and postpartum^4,5^, implicating both estrogen and progesterone in neuroplasticity driven by reproductive transitions in women. Importantly, these periods of significant hormonal shifts are associated with the increased symptomology and risk of mental disorders, including postpartum depression^6^ (affecting 20% of pregnant people), premenstrual dysphoric disorder^7^ (affecting 5-8% of menstruating individuals), and premenstrual exacerbation of mood disorders^8^ (occurring in >50% of affected women). Thus, it is crucial to characterize brain changes during these periods not only structurally, but also at cellular and molecular levels in order to gain molecular insights into brain changes and potentially find novel therapeutic targets.

Previous studies in mice focused on bulk neuronal (NeuN+) cells in the ventral hippocampus (vHIP) and found changes in chromatin organization and gene expression across the estrous cycle^9^. These chromatin dynamics were linked to changes in hippocampal dendritic spines and changes in anxiety-and depression-related behaviors, providing insights into hormone-driven structural plasticity and psychiatric risk^9^. Recently, a single cell study showed extensive hormone-driven gene expression and cellular state changes in the reproductive organs, providing a high resolution view of tissue remodeling that occurs across the estrous cycle^10^. In the brain, only one study used hormonal treatments with estradiol and progesterone in ovariectomized (OVX) rodents to mimic the estrous cycle stages and showed important hormone status-dependent changes in the estrogen receptor-rich hypothalamus^11^.

However, high-resolution cellular and molecular studies of the effects of physiological ovarian hormone shifts in the brain are missing. Here, we performed multiome (gene expression and chromatin accessibility) analysis of the mouse vHIP across sex, estrous cycle, and peripartum period at single cell resolution and revealed biological roles of dynamic cellular and molecular changes induced by ovarian hormone shifts.

## Cellular composition of the ventral hippocampus across the estrous cycle and sex

We focused on the vHIP due to its critical role in emotion and stress regulation in rodents^12^, and findings from rodents^1,9,13^ and humans^3,14,15^ showing that the hippocampus is particularly responsive to ovarian hormone shifts. A previous study characterized the cellular composition of the vHIP of male CD-1 (ICR) mice using single-cell RNA sequencing technology^16^. Here, we leveraged the single-nucleus multiome analysis of gene expression (RNA-seq) and chromatin accessibility (ATAC-seq) of >48,000 vHIP nuclei. We included C57BL/6J male mice as well as females of two estrous cycle stages – proestrus (high estradiol, low progesterone) and diestrus (low estradiol, high progesterone) – in order to refine the cellular classification of the vHIP across sexes and ovarian hormone shifts (**Figure 1A**). After integrating RNA and ATAC data, we performed a Weighted Nearest Neighbor analysis (WNN) at 0.8 resolution which identified 60 distinct cellular clusters in the vHIP (**Figure 1B, Extended Data Figure 1A**). Using known cell type marker genes, we confirmed the presence of eight broad cell types including excitatory (glutamatergic) neurons (70.2%), inhibitory (GABAergic) neurons (13.1%), astrocytes (7.3%), oligodendrocytes (3%), microglia (2.4%), oligodendrocyte precursor cells (2.2%), endothelial cells (1.4%), and choroid plexus cells (0.4%; **Figure 1C, Extended Data Figure 1B, 2A**). Using the MapMyCells platform^17^ and additional known marker genes (**Extended Data Figure 2B, Extended Data Table 1**), we performed further, semi-broad classification of vHIP excitatory (CA1, CA2, CA3, and dentate gyrus - DG) and inhibitory (Pvalb, Sst, Vip, Reln, Nos1, Calb2, and Cck) neurons (**Figure 1D**). We also identified a small population of excitatory neurons originating from other areas: entorhinal cortex (ENT), prosubiculum/presubiculum/parasubiculum (PPP), and subiculum (SUB) which were included in the semi-broad classification (**Figure 1D, Extended Data Fig. 1C**). All three cellular classifications (broad, semi-broad, and 60-cell subtypes) allowed us to ask different biological questions and provided different insights into the effect of ovarian hormone shifts and sex on the brain. For instance, at the 60-cell cluster level, we performed the analysis of cellular proportions and revealed remarkable dynamism of the vHIP cellular composition driven by the estrous cycle, affecting 11 different subpopulations of mainly excitatory neurons (**Figure 1E, Extended Data Figure 3A-B**). In addition, the effect of sex on cellular proportions in vHIP was found to be dynamic and dependent on the estrous cycle stage (**Figure 1E, Extended Data Figure 3C-D**), with larger sex effects found when females are in the high estrogenic, proestrus stage (17 different clusters) than in the low estrogenic diestrus state (10 clusters), indicating the role of estrogen’s shifts in driving cell state changes in the vHIP. We needed to use the highest resolution cell map of the vHIP (**Fig. 1B**) to detect cell proportion changes, which are likely mainly changes in the cell states rather than cell type^18^, since we have not observed these changes at the broad or semi-broad cell level.

**Figure 1.**
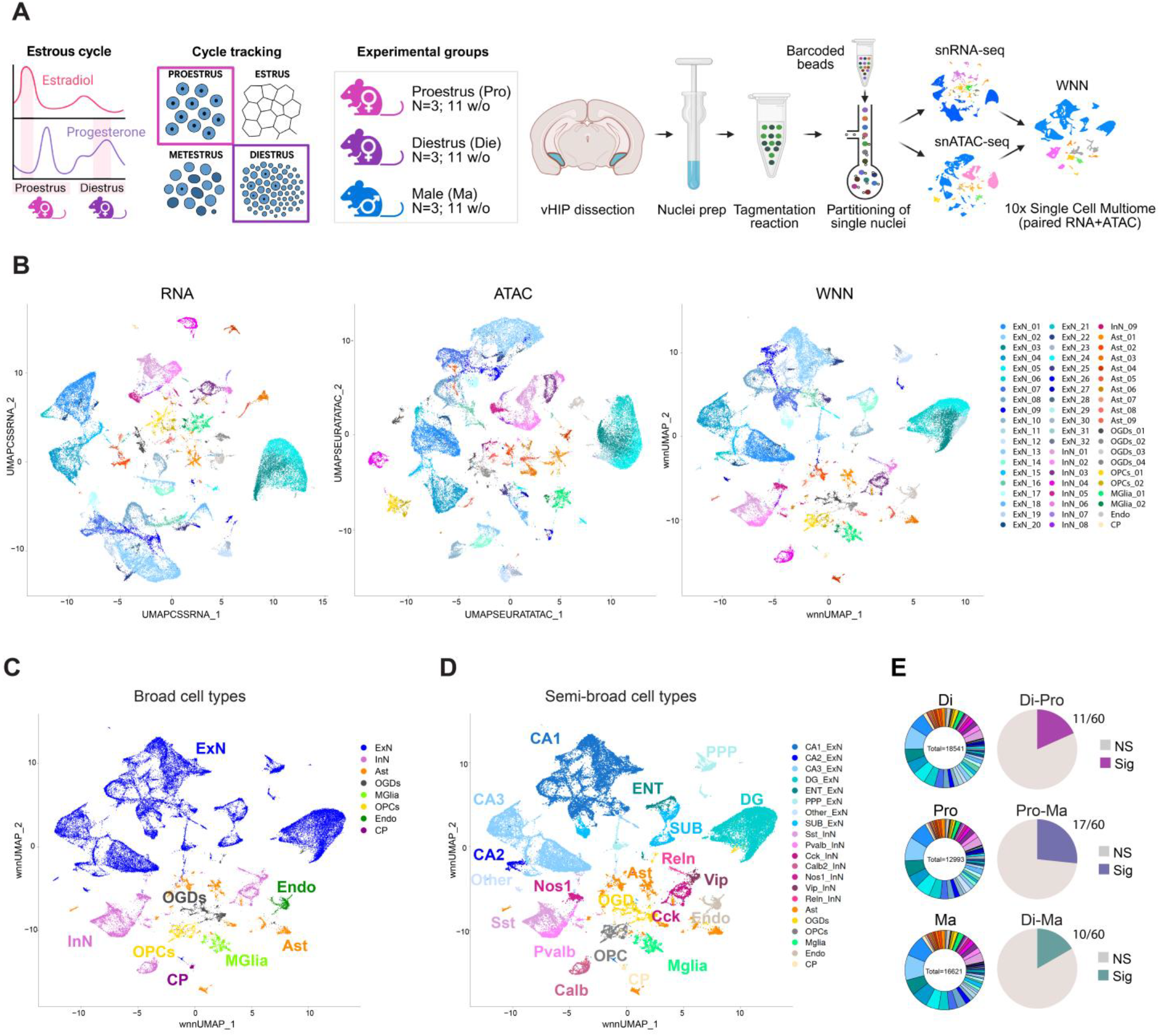
Single-nucleus multiome profiling reveals cell-type-resolved map of the ventral hippocampus across the estrous cycle and sex. **A**. Schematic of the mouse estrous cycle showing fluctuations in estradiol and progesterone across proestrus (Pro) and diestrus (Die). Female mice were staged by daily vaginal cytology for three consecutive cycles and classified as Pro or Die; age-matched males (Ma) served as comparison controls (N = 3 biological replicates or 6 animals/group). Ventral hippocampi (vHIP) were collected at 11 weeks of age and nuclei were isolated for 10x Genomics single-nucleus multiome (RNA-seq + ATAC-seq) assay. **B**. UMAP embeddings from single-nucleus gene expression (RNA), chromatin accessibility (ATAC), and weighted-nearest-neighbour (WNN) integration reveal consistent clustering of major neuronal and non-neuronal populations, with WNN providing enhanced resolution. **C**. Broad cell-type annotations identify excitatory (ExN) and inhibitory (InN) neurons, astrocytes (Ast), oligodendrocytes (OGDs), microglia (MGlia), oligodendrocyte precursor cells (OPCs), endothelial cells (Endo), and choroid plexus (CP) cells. **D**. Semi-broad annotations further resolve excitatory neuronal subtypes (CA1, CA2, CA3, DG, ENT, PPP, and SUB) and inhibitory neuron subclasses (Sst, Pvalb, Cck, Calb2, Nos1, Vip, and Reln). **E**. Cellular proportion analysis of the 60-cluster map (donut graphs, left) shows significant estrous cycle-and sex-dependent differences in vHIP cellular composition including 11 out of 60 clusters in Di-Pro comparison, 17 clusters in Pro-Ma comparison, and 10 clusters in Di-Ma comparison (two-sided proportion test, padj < 0.05; pie charts, right). DG, dentate gyrus; SUB, subiculum; ENT, entorhinal cortex; PPP, prosubiculum/presubiculum/parasubiculum; Sst, somatostatin; Pvalb, parvalbumin; Cck, cholecystokinin; Calb2, calbindin 2; Nos1, nitric oxide synthase 1; Vip, vasoactive intestinal peptide; Reln, reelin. Schematics were created with BioRender.

## Excitatory neurons are the major cell type in the vHIP affected by the estrous cycle and sex

To identify the effects of the estrous cycle and sex on gene expression and chromatin accessibility in the vHIP, we first performed analyses across broad cell types. We found the highest number of differentially expressed genes (DEGs) in excitatory neurons with very few DEGs in other cell types (**Extended Data Fig. 4A**). Importantly, the highest number of DEGs was found in the Diestrus-Proestrus comparison (N=1144) and DEGs were largely specific to each group comparison, with the highest overlap (N=399) found between Diestrus-Male and Proestrus-Male comparisons constituting sex-specific DEGs (**Extended Data Fig. 4B**). Enrichment analysis of sex-specific DEGs (**Extended Data Fig. 4C**) had shared terms with Diestrus-Proestrus comparison (**Extended Data Fig. 4D**) related to neurogenesis, synaptic organization, and dendritic spines. Importantly, we also found the enrichment of the Egr1 motif linked to estrous cycle-dependent (but not sex-specific) DEGs, indicating the role of the immediate early gene Egr1 in estrous cycle-dependent gene regulation which we reported previously^9,19^. Similar to previous study on bulk neuronal cells^9^, we also found that chromatin changes hugely exceeded gene expression differences both across the estrous cycle and sex (**Extended Data Fig. 4E**). Here, however, we were able to show that chromatin changes are found not only in excitatory neurons but across cell types (**Extended Data Fig. 4E**). In excitatory neurons, 19% of DEGs had at least one differentially accessible region (DAR), indicating that chromatin accessibility changes, at least in part, mediate changes in gene expression. Together, the data on broad excitatory neurons largely reproduce the estrous cycle-and sex-dependent changes we found in neuronal (NeuN+) cells previously^9^ and single cell data allow for more precise insights into not only neuronal cells but also across cell types including glial cells.

### Cell type-specific changes in gene expression in vHIP across the estrous cycle and sex

Our next step was to look into the semi-broad cell classification (**Fig. 1D**). Importantly, among excitatory neurons, we were able to identify a CA2 cell population, which previous RNA-seq studies reported as missing in the vHIP subregion^16,20^, possibly because the chromatin accessibility information was not available. However, while we had sensitivity to detect this small population of cells (0.9%, **Extended Data Fig. 1C**), we found very few DEGs and DARs in this region. We, therefore, focused on CA1, CA3, and DG excitatory neurons which, consistent with the broad cell data, exhibited the highest number of DEGs among all cell types (**Fig. 2A**). Importantly, unlike DEGs, chromatin changes were found across semi-broad cell types, including in inhibitory neurons and in glial cells, and were much more extensive in the Diestrus-Proestrus and Proestrus-Male comparison than in the Diestrus-Male comparison (**Fig. 2B**), implying that both within-and between-sex differences are largely driven by high estradiol levels in females. Overall, the highest number of DARs was found across the estrous cycle, particularly in CA1, CA3, and DG excitatory neurons (**Fig. 2B**).

**Figure 2.**
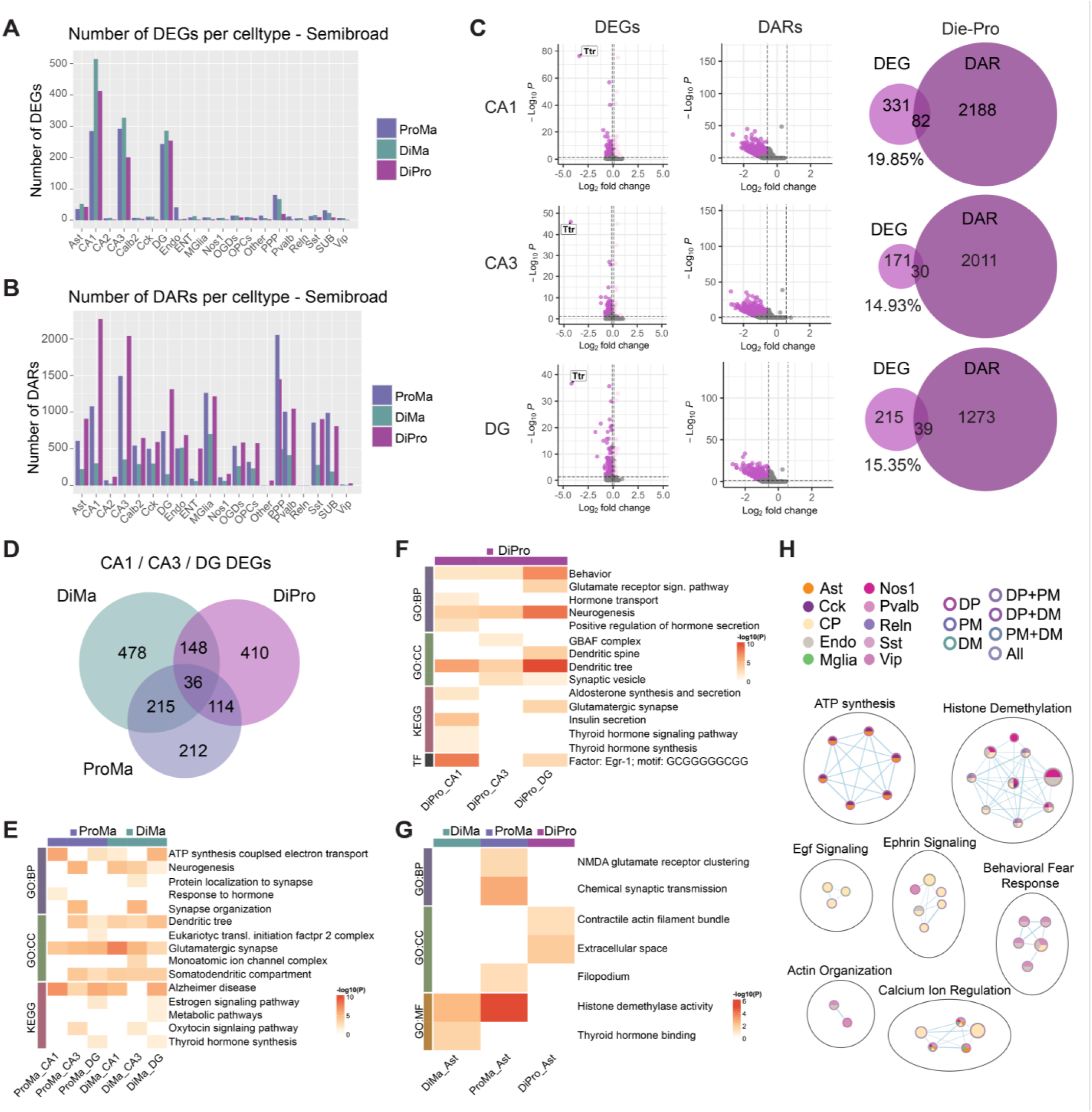
Gene expression and chromatin changes in semi-broad cell types of the ventral hippocampus. Bar graphs show the number of: **A**. differentially expressed genes (DEGs, padj < 0.05 & |log2fold-change| > 0.1), and **B**. differentially accessible regions (DARs, padj < 0.05 & |log2fold-change| > 0.58) across semi-broad cell types within proestrus-male (ProMa), diestrus-male (DiMa), and diestrus-proestrus (DiPro) group comparisons. **C**. Within DiPro comparison, volcano plots show DEGs (left, *Ttr* gene is highlighted) and DARs (middle), along with Venn diagrams showing the overlap between DEGs and genes annotated to DARs within the major excitatory neurons in the ventral hippocampus (vHIP): CA1 (top), CA3 (middle), and dentate gyrus (DG, bottom). **D**. A Venn diagram showing the overlaps across the three comparisons of DEGs pooled from CA1, CA3, and DG cell types. Heatmaps of enrichment analysis of: **E**. the 215 DEGs that overlap between ProMa and DiMa comparisons (i.e. sex difference DEGs; shown in (**D**)). **F**. all DiPro DEGs shown in (**D**) across CA1, CA3, and DG excitatory neurons. **G**. A heatmap showing the enrichment results of astrocyte DEGs across the three comparisons. **H**. An enrichment map depicting the results of GSEA analysis performed in inhibitory neurons and glial cells of the vHIP. Each node is an enriched gene set and the inner color of the node corresponds to cell type, while the color of the node’s border denotes the group comparison(s) in which the gene set is significant using the same color scheme as the Venn diagram in (**D**). Edges between nodes signify genes shared between gene sets with edge thickness being proportional to the number of shared genes. Enrichment heatmaps (**E, F, G**) show statistically significant terms and pathways (p_adj_ < 0.05) from Gene Ontology (GO) Biological Process (BP), Cellular Component (CC), or Molecular Function (MF) databases, as well as KEGG pathways and TRANSFAC motifs. Heatmap cells are colored by-log10(P_adj_) values. SUB, subiculum; ENT, entorhinal cortex; PPP, prosubiculum/presubiculum/parasubiculum; Sst, somatostatin; Pvalb, parvalbumin; Cck, cholecystokinin; Calb2, calbindin 2; Nos1, nitric oxide synthase 1; Vip, vasoactive intestinal peptide; Reln, reelin; oligodendrocytes (OGDs), microglia (MGlia), oligodendrocyte precursor cells (OPCs), endothelial cells (Endo).

Within each major excitatory neuron subtype (CA1, CA3, and DG), we found 201-413 DEGs and 1312-2270 DARs across the estrous cycle, with 15-20% gene overlap between DEGs and DARs, implying that chromatin accessibility changes drive gene expression differences, at least in part (**Fig. 2C**). Notably, overlap between DEGs and DARs was generally higher in the Diestrus-Proestrus comparison than it was in Male-Female comparisons (**Extended Data Fig. 5**). Among Diestrus-Proestrus DEGs, the *Ttr* gene, encoding a transporter protein Transthyretin, stood out in terms of fold change (>10.4 fold) and significance (FDR-adjusted p-value (Padj) < 2×10^−23^, **Extended Data Table 2**) and was common to all major subfield excitatory neuron subtypes – CA1, CA3, and DG (noting that some excitatory neuron subtypes showed undetectable *Ttr* expression in Diestrus**; Fig. 2C**).

While our overall number of unique DEGs in semi-broad excitatory neurons was somewhat lower compared to broad excitatory neurons (**Extended Data Fig. 4G**), we observed more specific enrichment terms when analyzing semi-broad cell types, likely reflecting more specific cellular classification. Again, we found that the majority of DEGs were specific to each group comparison (**Fig. 2D**). General sex-specific genes (N=215), which overlapped between Diestrus-Male and Proestrus-Male comparisons, shared terms such as neurogenesis, glutamatergic synapse and dendritic tree across excitatory neuronal subtypes (**Fig. 2E)**, which were found in broad excitatory neurons as well (**Extended Data Fig. 4C**). However, in the semi-broad analysis, we also found terms that were specific to vHIP subregions, such as oxytocin signaling pathway (unique to CA3), as well as estrogen signaling, metabolic pathways, and thyroid hormone synthesis (unique to DG; **Fig. 2E)**, which were not previously found in the broad cell analysis (**Extended Data Fig. 4C**). Focusing on the estrous cycle DEGs (N=708, **Fig. 2D**), we further found terms such as thyroid hormone signaling (specific to CA1), chromatin remodeling GBAF complex (specific to CA3), and dendritic spine (specific to DG, **Fig. 2F**). It is also interesting to note that terms such as behavior, neurogenesis, and dendritic tree showed much stronger enrichment in DG compared to CA1 and CA3, which is particularly important for neurogenesis which is a DG-specific process (**Fig 2F**). We again found enrichment of the Egr1 motif in the estrous cycle DEGs, but only in the CA1 (strong) and DG (moderate, **Fig 2F**).

Beyond excitatory neurons, only astrocytes had a sufficient number of DEGs to perform the robust Gene Ontology (GO) enrichment analysis (e.g. in the estrous cycle comparison, N=42), which interestingly found the enrichment of the term Extracellular space in the Diestrus-Proestrus comparison only, consistent with astrocytes’ role in regulating the extracellular concentration of glutamate (**Fig. 2G**). While few DEGs were identified outside core excitatory clusters, we performed Gene Set Enrichment Analysis (GSEA) on lists of ranked gene expression differences from inhibitory neurons and glial populations to determine whether functionally relevant shifts in gene set expressions could be identified. We find enrichment for genes related to: i) ATP synthesis in Astrocytes and Cck neurons; ii) Egf and Ephrin signaling genes in choroid plexus cells; iii) behavioral fear response enrichment in endothelial and Pvalb cells; and a more general enrichment for calcium regulation and chromatin regulation across cell types (**Fig. 2H**).

### Cell type-specific changes in chromatin accessibility in vHIP across the estrous cycle and sex

The finding of more extensive chromatin than gene expression changes within excitatory neuronal subtypes, as well as across all cell types, is of great interest, as it implies that chromatin changes may have effects that go beyond immediate changes in gene transcription. Interestingly, we identified more unique DARs in semi-broad cell types than we did in broad excitatory neurons (**Extended Data Fig. 4H**). We then looked at the chromatin changes in between-group comparison across excitatory neurons, inhibitory neurons, and glial cells (**Fig. 3A-C**), and observed a similar pattern: unlike what we find with DEGs where the biggest overlap is in sex-specific comparisons (**Fig. 2D**), here the biggest overlap is between Diestrus-Proestrus and Proestrus-Male comparisons across cell types (**Fig. 3A-C**), indicating that estrogen levels are driving chromatin differences across the cycle and between the sexes in all vHIP cell types.

**Figure 3.**
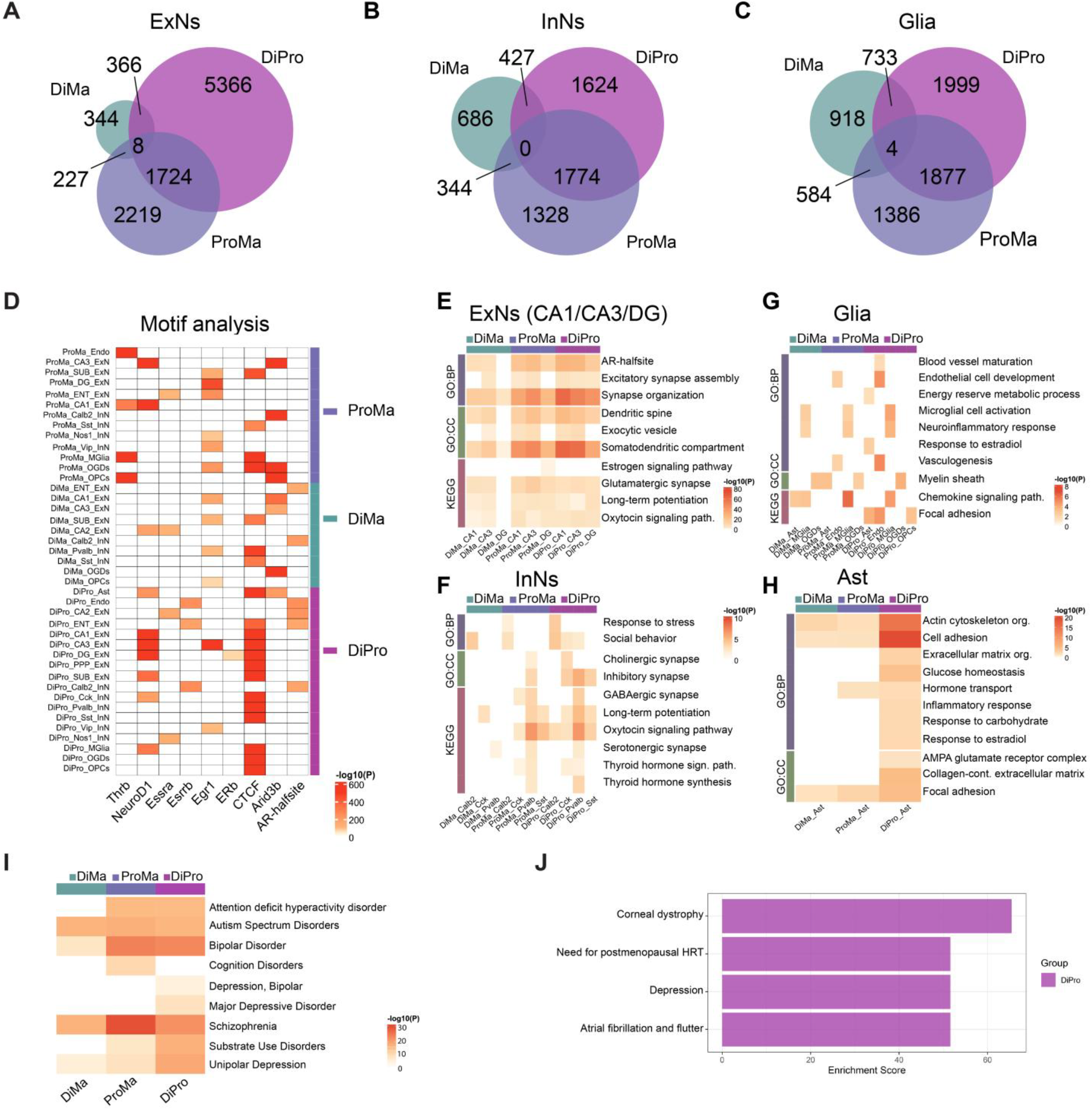
Cell type-specific chromatin priming in the ventral hippocampus across the estrous cycle and sex. Venn diagrams show overlapping differentially accessible regions (DARs, padj < 0.05 & |log2fold-change| > 0.58) in the Diestrus-Proestrus (DiPro), Diestrus-Male (DiMa), and Proestrus-Male (ProMa) comparisons across excitatory neurons (ExNs, **A**), inhibitory neurons (InNs, **B**), and glial cells (Glia, **C**) of the ventral hippocampus. **D**. Motif analysis depicts both the top 3 (by maximum −log10p) and selected (by relevance to hormonal signaling) transcription factor motifs that are enriched in DARs across all semi-broad cell types and group comparisons. Enrichment analysis of genes annotated to DARs is shown for: **E**. major ventral hippocampal ExNs (in CA1, CA3, and DG); **F**. InNs; **G**. Glial cells (all), and **H**. astrocytes (Ast). Enrichment heatmaps (**E, F, G, H**) show statistically significant terms and pathways (p_adj_ < 0.05) from Gene Ontology (GO) Biological Process (BP) and Cellular Component (CC) databases, as well as KEGG pathways. **I**. A heatmap showing the enrichment of genes annotated to DARs for genetic associations with human brain disorder (assessed using the DisGeNET database^25^) in all three group comparisons. **J**. A bargraph depicting the enrichment of DiePro-specific DAR-annotated genes for human disorders (assessed using the PheWeb database^26^; padj < 0.05), with enrichment score calculated as the log(P) of the Fisher’s exact test multiplied by the z-score of the deviation from the expected rank. Heatmap cells are colored by −log10(P_adj_) values. DG, dentate gyrus; SUB, subiculum; ENT, entorhinal cortex; PPP, prosubiculum/presubiculum/parasubiculum; Sst, somatostatin; Pvalb, parvalbumin; Cck, cholecystokinin; Calb2, calbindin 2; Nos1, nitric oxide synthase 1; Vip, vasoactive intestinal peptide; Reln, reelin; oligodendrocytes (OGDs), microglia (MGlia), oligodendrocyte precursor cells (OPCs), endothelial cells (Endo).

Within chromatin accessibility changes, we found the enrichments of multiple motifs that are either generally important for chromatin organization (CTCF and Arid3b) or directly relevant to hormone-mediated regulation (Egr1, estrogen response elements [EREs], Estrogen-Related Receptor Elements [ERREs], etc.; **Fig. 3D**). The enrichment of the CTCF motif was particularly strong for the estrous cycle-dependent chromatin changes, across cell types (**Fig. 3D**), which is consistent with our previous finding of extensive 3D genome changes across the estrous cycle^21^. We found the enrichment of EREs in DG of the Diestrus-Proestrus comparison only, but this was not surprising as we previously observed a similar phenomenon in our bulk NeuN+ cells^9^, where there was Egr1 (but not ERE) enrichment in chromatin accessibility data across the estrous cycle. Another motif that showed the estrous cycle-specific enrichment in the current study was NeuroD1, including in CA1, CA3, and DG excitatory neurons, as well as in Cck inhibitory neurons and microglia (**Fig. 3D**). Importantly, for the NeuroD1 finding, we needed to have the analysis of the semi-broad cell types, since this pioneer factor^22^ showed no enrichment in broad neuron cell types in this study (**Extended Data Fig. 4I**) or in our previous study on NeuN+ cells^9^. Overall, our data indicate that ovarian hormones may mediate chromatin reorganization through cell type-specific recruitment of pioneer factors such as Egr1^19^ and NeuroD1^22^, in interaction with 3D chromatin changes likely mediated via CTCF and estrogen receptors^21,23^.

We next performed enrichment analysis on chromatin accessibility data across cell types in order to examine the functional relevance of chromatin changes (**Fig. 3E-H**). Excitatory neurons (**Fig. 3E**) showed a lot of terms that mirrored enrichment analysis of gene expression data (**Fig. 2E-F**), and were also consistent with enrichment analysis of the genes that showed changes in both chromatin and gene expression (**Extended Data Fig. 6**). These enrichment terms included synapse organization, dendritic spines, and glutamatergic synapse (**Fig. 3E**), confirming that excitatory neurons in CA1, CA3, and DG subregions mediate changes in brain function and behavior across the estrous cycle. Notably, in inhibitory neurons (**Fig. 3F**), chromatin accessibility data showed enrichment of multiple neurotransmitter systems (GABAergic, cholinergic, and serotonergic), hormone signaling pathways (oxytocin and thyroid signaling pathways), and response to stress, which were cell subtype specific and often primarily enriched in the estrous cycle-dependent manner (**Fig. 3F**). Within glial cells, we found many interesting cell type-specific enrichment terms including: endothelial cell development, local adhesion, and vasculogenesis for endothelial cells; microglial activation and neuroinflammatory response for microglia; and myelin sheath for oligodendrocytes (**Fig. 3G**). While some of these terms were both estrous cycle-and sex-dependent, in astrocytes, more specifically, we find many terms that are estrous cycle-specific including extracellular matrix organization, glucose homeostasis, response to estradiol, and AMPA glutamate receptor complex (**Fig. 3H**). These data indicate that, while many of the chromatin changes in inhibitory neurons and glial cells are not converted to gene expression changes during the physiological estrous cycle, they have the potential to affect the genes that are both related to their specific cellular function or to the function of neighboring neuronal cells if the need for this arises such as during pregnancy or stressful events.

To address the possibility that chromatin changes during the cycle may also predispose the brain to stress-sensitive disorders, we used the EnrichR platform^24^ to assess whether genes annotated to excitatory neuron (CA1, CA3, and DG) DARs had genetic associations with brain disorders in humans. Strikingly, querying both the DisGeNET^25^ (**Fig. 3I**) and PheWeb^26^ (**Fig. 3J**) databases indicated that genes annotated to estrous cycle-dependent DARs show enrichment for several neuropsychiatric disorders in humans, with particularly prominent enrichment for depression across the two databases (**Fig. 3I-J**).

## Dynamic cellular changes in the dentate gyrus across the estrous cycle

While, so far, DG showed unique features in terms of gene expression and chromatin changes compared to CA1 and CA3 excitatory neurons (**Fig. 2-3**), our highest resolution 60-cell cluster vHIP map revealed 5 different cellular subtypes in DG (**Fig. 4A**). This was in contrast to the maps we generated when we uploaded our data into the MapMyCell platform^17^, which revealed only one DG cluster even at the resolution where 126 clusters were detected (**Fig. 4A**). Our increased precision in DG cell classification further emphasizes the power of having single cell chromatin accessibility data, alongside gene expression, to call cellular clusters. Importantly, based on gene markers (**Extended Data Table 1**), we were able to classify 4 of the 5 DG cell clusters into the following biologically relevant cell subtypes: neural stem progenitor cells (NSPC), immature granule neurons (ImGN), mature granule neurons (MGN), and Penk^+^ mossy cells (**Fig. 4A**). These cell subtypes are consistent with the neurogenesis process happening in the DG^27^. We next made a comparison of the four DG cell subtypes based on their gene expression and chromatin accessibility profiles (**Fig. 4B**). In this comparison, NSPCs showed the most open, flexible chromatin state consistent with their role as nondifferentiated neural stem cells (**Fig. 4B**). In contrast, other cell subtypes showed more distinct chromatin and transcriptional states (**Fig. 4B**), consistent with their more advanced differentiated cellular state.

**Figure 4.**
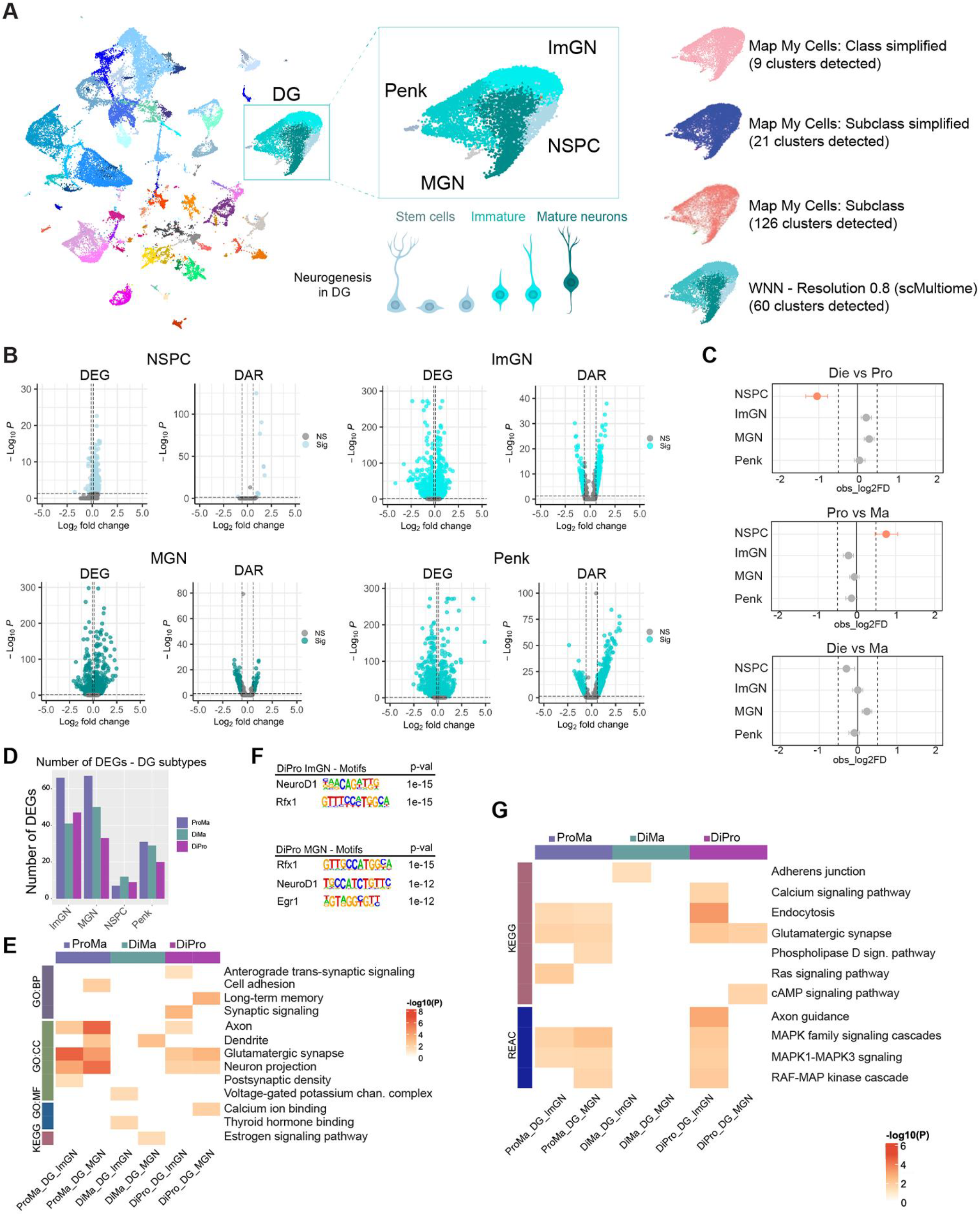
High-resolution multiome profiling reveals estrous cycle-and sex-dependent shifts in dentate gyrus neurogenic-lineage composition. **A**. UMAP visualization of the weighted-nearest-neighbor (WNN)-integrated dataset highlighting the dentate gyrus (DG) cluster (Left). The inset shows progressive refinement of DG with increasing clustering resolution including four transcriptionally and epigenetically distinct DG populations: mature granule neurons (MGN), immature granule neurons (ImGN), neural stem/progenitor cells (NSPC), and Penk-expressing mossy cells (Penk; top-middle). Colors correspond to WNN-derived subclusters and are consistent across panels including in the scheme of the neurogenesis process in the DG showing stem cells, immature and mature neurons (bottom-middle). Automated reference-based annotation (MapMyCells) failed to distinguish these subtypes, which emerged only through manual high-resolution analysis (Right). **B**. Volcano plots showing differentially expressed genes (DEGs, p_adj_ < 0.05 & |log_2_fold-change| > 0.1; left) and differentially accessible regions (DARs, p_adj_ < 0.05 & |log_2_fold-change| > 0.58); right) for each DG subcluster using one-vs-rest comparisons. Blue points painted based on the DG cell cluster color code indicate significant genes/regions. **C**. Cell proportion analysis reveals group-dependent differences in the proportion of DG subclusters, including a significant reduction in NSPC proportion in diestrus relative to proestrus and in proestrus relative to males (two-sided proportion test, padj < 0.05). **D**. Bar graphs show the number of: DEGs (p_adj_ < 0.05 & |log_2_fold-change| > 0.1) across DG subclusters within proestrus-male (ProMa), diestrus-male (DiMa), and diestrus-proestrus (DiPro) group comparisons. **E**. A heatmap shows the enrichment analysis of DEGs in ImGN and MGN cell clusters in all three group comparisons. **F**. Motif analysis of chromatin changes in ImGN (top) and MGN (bottom) cells in the Di-Pro comparison (shown are the results of the top three motifs; note: ImGN analysis retrieved only 2 significant motifs total). **G**. A heatmap shows the enrichment analysis of DARs (p_adj_ < 0.05 & |log_2_fold-change| > 0.58) across ImGN and MGN cell clusters in all three group comparisons. Enrichment heatmap shows statistically significant pathways (p_adj_ < 0.05) from KEGG and Reactome (REAC) pathway analysis. Heatmap cells are colored by −log10(P_adj_) value.

We next explored the cell proportion data at the level of the 60-cell cluster map (**Fig. 1E, Extended Data Fig. 3**) but now focusing on the four DG cell subtypes (**Fig. 4C**), exploring the possibility that the estrous cycle and sex may not only affect cell states but also cell (sub)types in the vHIP. Importantly, we found that the NSPC population shows particular dynamism with diestrus mice exhibiting a higher proportion of this cell type compared to proestrus females (**Fig. 4C**). Since previous studies showed that estrogen increases neurogenesis during proestrus^28^, our finding is consistent with the idea that NSPCs are utilized during proestrus to make new neurons while they are being replenished during the low estrogenic, diestrus phase. We also find that males have a higher proportion of NSPCs compared to proestrus, but not diestrus females, further revealing another dynamic, functionally-relevant sex difference (**Fig. 4C**).

Since only NSPCs show changes in cell proportions, we were interested in exploring changes in gene expression and chromatin in other DG cell subtypes across the estrous cycle and sex (**Fig. 4D-G**). The highest number of DEGs was found in immature and mature granule neurons (**Fig. 4D**), and given their importance for neurogenesis and vHIP function, we focused on these two cell types. We again found many terms relevant for neuronal function such as glutamatergic synapse, neuron projection, and axon, but these terms appeared to be shared between Diestrus-Proestrus and Proestrus-Male comparison, but not Diestrus-Male comparison (**Fig. 4E**). This further implies that cellular proportion changes in NSPCs and transcriptional changes in ImGN and MGN cells in Die-Pro and Pro-Male comparison support the neurogenesis process driven by increased estrogen in proestrus.

It is of interest to note that motif analysis of the chromatin accessibility data revealed that two transcription factors involved in neurogenesis – NeuroD1^29^ and Rfx1^30^ – may be driving chromatin changes in both ImGN and MGN cells across the estrous cycle (**Fig. 4F**). Interestingly, only mature neurons show the enrichment of the immediate early gene product Egr1 (**Fig. 4F**). These motifs were specific to the estrous cycle and were not found among the top motifs in the female-male comparisons (**Extended Data Fig. 7**). Further pathway analysis of chromatin changes showed that multiple kinase pathways, including the MAPK family signaling and the RAF-MAP kinase cascade may be involved downstream of estrogen in changing chromatin and gene expression in immature (but not mature) neurons during the high-estrogenic stage of the cycle (**Fig. 4G**).

### Peripartum changes in the vHIP cells partially mimic changes across the estrous cycle

The estrous cycle involves systemic fluctuating ovarian hormone shifts across 4-5 days in mice (**Fig. 1A)**. To further explore the role of ovarian hormone shifts on the cellular and molecular make-up of the vHIP, we performed another multiome study which compared two groups with a more extreme ovarian hormone shift: mice in late pregnancy (gestational day - GD18), which have high levels of estradiol and progesterone, with mice within 24 hour postpartum that experience a sudden withdrawal of both ovarian hormones^31,32^ (**Fig. 5A**). Once single cell gene expression and chromatin accessibility data were integrated, using >30,000 nuclei, we identified 46 cellular clusters (**Extended Data Fig. 8A**) and the same broad (**Extended Data Fig. 8B**) and semi-broad (**Fig. 5B**) cell types in the vHIP in our peripartum dataset. Importantly, we found that 10 out of 46 cell clusters in the vHIP showed changes in cellular proportion in pregnancy when compared to the vHIP postpartum (**Extended Data Fig. 8C**). Remarkably, similar to what we found during the estrous cycle (**Fig. 4C**), the NPSC population in the DG showed changes from pregnancy to postpartum, with higher proportion found during pregnancy (**Fig. 5C**), confirming the sensitivity of this cellular population to dynamic ovarian hormone shifts.

**Figure 5.**
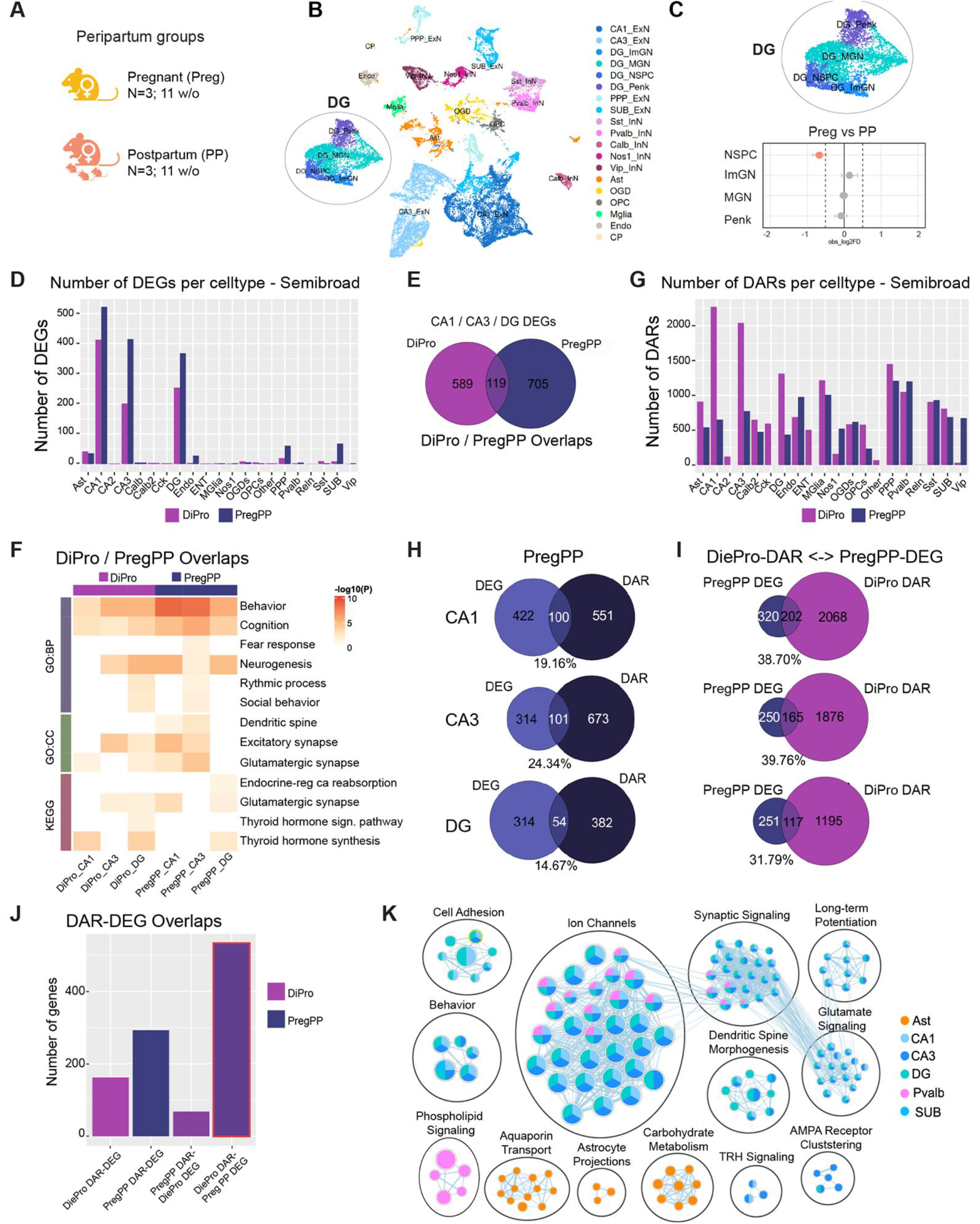
Comparison of the estrous cycle and peripartum single cell profiles in the ventral hippocampus. **A**. Schematic of the peripartum groups of pregnant (Preg) and postpartum (PP) mice (N = 3 biological replicates or 3 animals/group). Ventral hippocampi (vHIP) were collected at 11 weeks of age, and nuclei were isolated for 10x Genomics single-nucleus multiome (RNA-seq + ATAC-seq) analysis. **B**. UMAP of weighted-nearest-neighbor (WNN) integration reveals consistent clustering of semi-broad cell populations in the vHIP. **C**. Focusing on dentate gyrus (DG) subregion, four cell subtypes are found including neural stem progenitor cells (NSPC), immature granule neurons (ImGN), mature granule neurons (MGN), and Penk+ cells. Cell proportion analysis reveals a decrease in the proportion of NSPCs in the postpartum group relative to pregnant mice (two-sided proportion test, padj < 0.05). **D**. A bar graph showing the number of differentially expressed genes (DEGs; p_adj_ < 0.05 & |log_2_fold-change| > 0.1) detected across semi-broad cell types in the diestrus-proestrus (DiPro, purple) and pregnancy-postpartum (PregPP, dark blue) group comparisons. **E**. A Venn diagram showing the overlap between datasets of DEGs in DiPro and PregPP groups pooled from the major vHIP excitatory neuron subtypes: CA1, CA3, and DG. **F**. A heatmap showing the enrichment results for the 119 DEGs that overlap the two datasets in (**E**). The heatmap contains statistically significant terms and pathways (p_adj_ < 0.05) from Gene Ontology (GO) Biological Process (BP) and Cellular Component (CC), as well as KEGG pathways; heatmap cells are colored by −log10(P_adj_) values. **G**. A bar graph showing the number of differentially accessible regions (DARs; p_adj_ < 0.05 & |log_2_fold-change| > 0.58) across semi-broad cell types in the DiPro and PregPP group comparisons. **H**. Overlap between DEGs and genes annotated to DARs in the PregPP group comparison in CA1 (top), CA3 (middle), and DG (bottom) excitatory neurons. **I**. An overlap of genes annotated to DiePro DARs with PregPP DEGs in CA1 (top), CA3 (middle), and DG (bottom) excitatory neurons. **J**. A bar graph demonstrating that the total number of DiePro DAR – PregPP DEG overlaps across all cell types exceeds the other possible DAR-DEG overlaps both within and across datasets. **K**. An enrichment map of results based on the DiePro DAR – PregPP DEG overlaps (bar outlined in red in **J**). Each node is a significant gene set and the inner color of the node corresponds to cell type. Edges between nodes denote genes shared across gene sets and the thickness of the edge is proportional to the number of shared genes. SUB, subiculum; ENT, entorhinal cortex; PPP, prosubiculum/presubiculum/parasubiculum; Sst, somatostatin; Pvalb, parvalbumin; Cck, cholecystokinin; Calb2, calbindin 2; Nos1, nitric oxide synthase 1; Vip, vasoactive intestinal peptide; Reln, reelin; Penk, Penk-expressing mossy cell; oligodendrocytes (OGDs); microglia (MGlia); oligodendrocyte precursor cells (OPCs); endothelial cells (Endo).

We next examined DEGs and DARs during the peripartum transition (**Extended Data Fig. 9A**) and compared them to the estrous cycle dataset (**Fig. 5D-K, Extended Data Fig. 9B**). Similar to the estrous cycle gene expression data, the majority of peripartum DEGs are found in vHIP subfield excitatory neuron populations CA1, CA3, and DG (**Fig. 5D**). However, we identified a higher number of DEGs in the peripartum dataset, particularly in CA3 neurons (**Fig. 5D, Extended Data Fig. 9A**), likely reflecting the greater ovarian hormone shift occurring during the pregnancy to postpartum transition. Comparing the excitatory neuron DEGs across the estrous cycle and peripartum shows that most DEGs are hormone transition-specific (**Fig. 5E**). However, overlapping DEGs (N=119) are enriched for many terms related to behavior, cognition, neurogenesis, dendritic spines, and hormone signaling (**Fig. 5F**), indicating shared functional effects of hormonal fluctuations across the transitions. The analysis of the peripartum-specific genes (N=705; **Fig. 5E**), show unique enrichment of the hormonal – estrogen, oxytocin, and GnRH – signaling pathways as well as enhanced enrichment of neurogenesis-, dendrite-, and postsynapse organization-relevant genes in the peripartum compared to the estrous cycle dataset, with all of these processes obviously needing enhancement and amplification during pregnancy and postpartum transition (**Extended Data Fig. 9B**).

Chromatin accessibility changes also show similarity between the estrous cycle and peripartum. Similar to the Diestrus-Proestrus comparison, we find that peripartum chromatin changes are more extensive than peripartum gene expression changes, and they occur across cell types, not only in excitatory neurons (**Fig. 5G**). Remarkably, though, the estrous cycle-driven chromatin changes in the vHIP CA1, CA3, and DG neurons greatly exceed those found peripartum (**Fig. 5G**), which is exactly opposite from what we find with DEGs (**Fig. 5D**). Accordingly, we find more limited enrichment of the NeuroD1 and ERE motifs in chromatin accessibility data during the peripartum transition (**Extended Data Fig. 9C**). Enrichment of Egr1 and particularly CTCF motif was more similar across the two datasets (**Fig. 3D, Extended Data Fig. 9C**), again implying the coordination between pioneer factors and 3D genome organizers in chromatin regulation by ovarian hormones. Similar to the cycle, around 15-25% of peripartum DEGs show both changes in gene expression and in chromatin accessibility across CA1, CA3, and DG neurons (**Fig. 5H**), indicating that, at least in part, peripartum chromatin changes mediate changes in gene expression.

We next wanted to address whether chromatin changes seen across the estrous cycle may at least in part be preparatory, priming the genome for gene expression changes that may occur in the case of pregnancy. To address this question, we overlapped DEGs in the peripartum dataset with DARs in the estrous cycle dataset (**Fig. 5I**). Remarkably, in this case, the overlap between DEGs and DARs increased approximately two-fold across CA1 (from 19.2% to 38.7%), CA3 (from 24.3% to 39.8%) and DG (from 14.7% to 31.8%, **Fig. 5I**), indicating that chromatin changes across the cycle may indeed prime the genome for gene expression changes in the vHIP required for pregnancy. In terms of absolute gene numbers, when we overlapped DARs and DEGs within and across datasets, the number of overlapping genes went from <200 in the estrous cycle DAR-DEG overlap and ~300 in peripartum DAR-DEG overlap to >500 in the estrous cycle DAR - peripartum DEG overlap (**Fig. 5J**). The number of overlapping genes actually decreased in peripartum DAR – estrous cycle DEG overlap (**Fig. 4J**), as expected for two separate datasets, further supporting the specificity of the finding that estrous cycle chromatin is primed for gene expression changes that are only realized in the presence of an altered milieu, such as during the peripartum period.

To address the possible functional relevance of genes that are primed during the estrous cycle and show changes peripartum, we performed enrichment analysis on genes overlapping the estrous cycle-DARs and peripartum-DEGs (**Fig. 5K**). In excitatory neurons, we found terms related to glutamate signaling, LTP, dendritic spines, behavior, thyroid signaling, AMPA receptor clustering, and cell adhesion. In Pvalb neurons, we find enrichment for phospholipid signaling. In Astrocytes, we see enrichment for metabolism, aquaporin water transport (important for blood-brain-barrier maintenance), and astrocyte projections, all of which are likely to support altered brain demands during pregnancy and postpartum transitions.

Finally, given the dramatically increased risk for psychiatric disorders postpartum^6^, we tested whether genes annotated to excitatory neuron (CA1, CA3, and DG) DEGs or DARs in the peripartum dataset had genetic associations with brain disorders in humans (**Extended Data Fig. 10**). Intriguingly, using the DisGeNET^25^ platform, we identified significant enrichment of multiple neuropsychiatric disorders in both postpartum DEGs and DARs, including depression, bipolar disorder, and schizophrenia (**Extended Data Fig. 10**). However, the enrichment status was stronger for DEGs in this case, likely reflecting functionally increased risks for these disorders postpartum, through hormonally-driven changes in gene expression.

### The transthyretin gene responds to ovarian hormone shifts

Finally, we focused on the *Ttr* (transthyretin) gene that was one of the top DEGs both across the estrous cycle (**Fig. 2C, Fig. 6A, Extended Data Fig. 11A**) and peripartum (**Fig. 6B**). *Ttr* encodes the transporter for thyroxine and retinoic acid and was previously shown to be an estrogen responsive gene, containing an ERE in its putative regulatory region^33^. We observed a robust increase in *Ttr* expression in proestrus compared to diestrus and during pregnancy compared to postpartum across cell types (**Extended Data Table 2**), consistent with *Ttr*’s responsiveness to rising estrogen levels. Importantly, it was previously reported that hippocampal *Ttr* may represent the dissection contamination derived from the choroid plexus^34^. While we found the choroid plexus (CP) cluster that clearly had *Ttr* as a marker gene (**Extended Data Fig. 2A**), we were able to show that *Ttr* differences were found across neuronal and glial cell clusters but not in the CP cluster (**Extended Data Fig. 11B; Extended Data Table 2**). The *Ttr* levels were higher in Proestrus than in Diestrus group across replicates (**Extended Data Fig. 11C**), removing the possibility that our result may be a product of random dissection differences among the samples.

**Figure 6.**
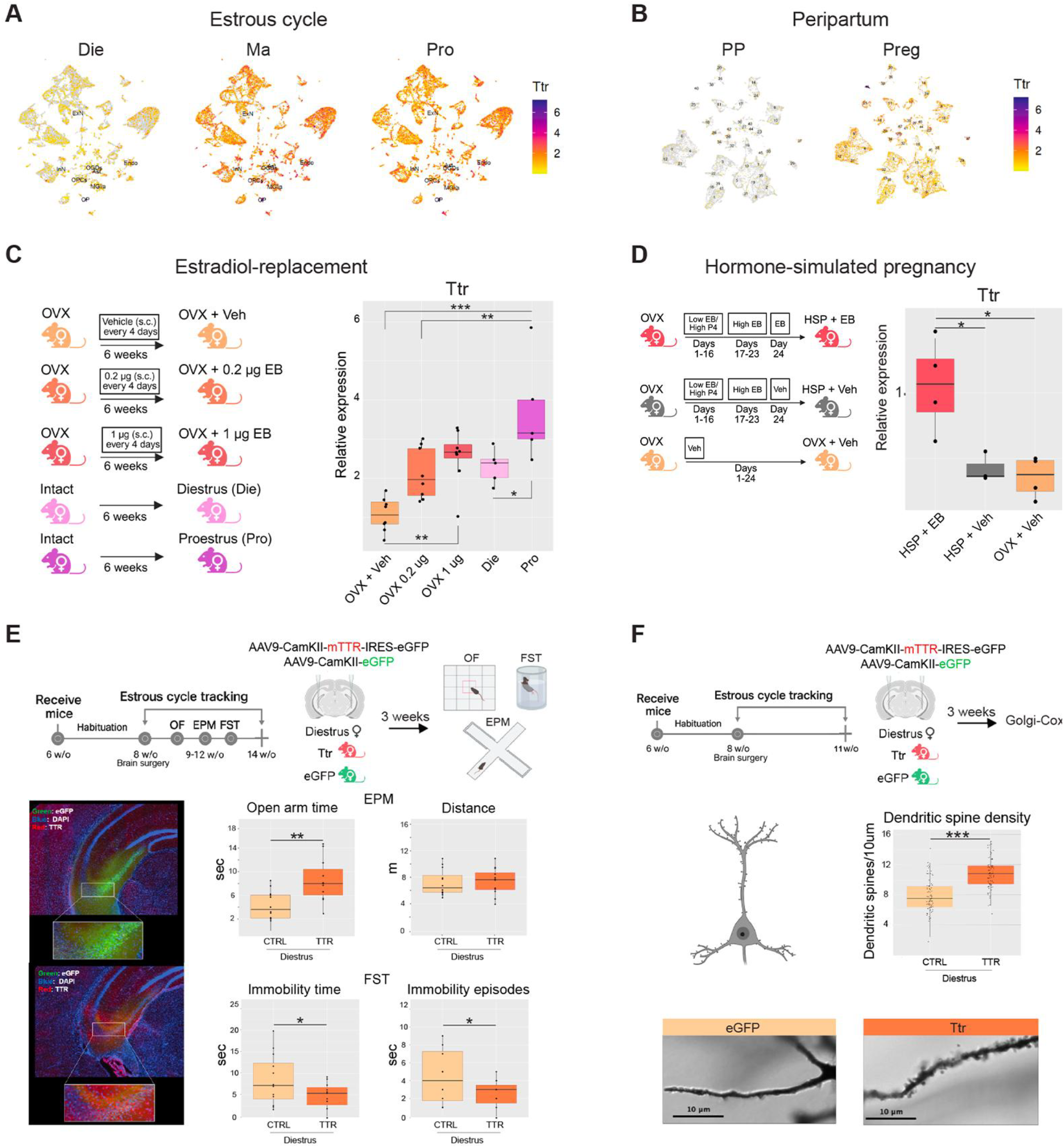
Ttr is hormonally regulated across the estrous cycle and peripartum states and functionally modulates ventral hippocampal structure and behavior. UMAP feature plots showing Ttr expression in: **A**. Diestrus (Die), Males (Ma), and Proestrus (Pro); and **B**. in pregnancy (Preg) and postpartum (PP). Color indicates average normalized expression. **C**. Experimental timeline for the cyclical estradiol-replacement paradigm (Left) and qRT-PCR analysis of Ttr expression (Right) in OVX females treated with either vehicle (OVX+Veh) or two different doses (0.2 µg and 1 µg; s.c.) of estradiol benzoate (EB), alongside proestrus (Pro) and diestrus (Die) cycling females (one-way ANOVA with Tukey post hoc test; *, p < 0.05; **, p < 0.01; ***, p < 0.001; N = 5-8 animals/group). **D**. Experimental timeline for the hormone-simulated pregnancy (HSP) paradigm (Left) and qRT-PCR analysis of Ttr expression in estrogen-sustained HSP animals (HSP + EB), estrogen-withdrawn HSP animals (HSP + Veh) and ovariectomized vehicle controls (OVX + Veh; one-way ANOVA with Tukey post hoc test; *, p < 0.05; N = 3-4 animals/group). HSP paradigm: Days 1-16: mice received daily s.c. injections of EB (0.5 μg) and progesterone (P4; 80 μg); Days 17-23: only EB was given at 10 μg/day; Day 24: mice either continued EB (HSP + EB) or were switched to vehicle (HSP+Veh) to model postpartum estrogen withdrawal. **E**. Experimental timeline for Ttr overexpression experiments and behavioral testing (top). Immunofluorescence confirms robust Ttr overexpression in vHIP excitatory neurons (bottom left; upper image, control; lower image, Ttr overexpression; green, eGFP; red, TTR; blue, DAPI). Box plots show open-arm time (one-tailed Mann-Whitney *U* test; **, p < 0.01) and total distance traveled (unpaired one-tailed *t*-test; p > 0.05) in the elevated plus maze (EPM), as well as immobility time and immobility episodes in the forced swim test (FST; unpaired one-tailed *t*-test; *, p < 0.05) in Ttr-overexpressing (Ttr) and control females (eGFP) in diestrus (bottom right; N = 10-12/group); **F**. Experimental timeline for Ttr overexpression experiments and Golgi-Cox staining (top). Box plots show dendritic spine density on CA1 pyramidal neurons in Ttr and eGFP females in diestrus (middle; unpaired one-tailed t-test, ***, p < 0.001; N = 5 animals/group). Representative photographs are shown for Ttr and eGFP females (10 µM scale). Box plots show the median (center line), 1st–3rd quartiles (box), and 1.5xIQR whiskers. Schematics created with BioRender.

To further provide the evidence that *Ttr* is hormonally regulated and to confirm our results from the single cell analysis, we used two hormone treatment paradigms – cyclical estrogen treatment that mimics estradiol changes across the estrous cycle^35^ (**Fig. 6C**) and the hormone simulated pregnancy (HSP) model that mimics estradiol and progesterone changes across pregnancy and postpartum^36,37^ (**Fig. 6D**). Importantly, using bulk qRT-PCR we were able to reproduce the increase in *Ttr* expression in the vHIP of proestrus compared to diestrus mice in a new animal cohort (**Fig. 6C;** one-way ANOVA, F(4,29)=10.47; p<0.001; Tukey’s post hoc test, Pro vs. Die, p=0.03). We were also able to show a dose-dependent increase in *Ttr* expression following cyclical estrogen treatment in OVX mice (**Fig. 6C;** Tukey’s post hoc test; OVX-Veh vs. OVX-1µg, p=0.003; OVX-Veh vs. OVX-0.2µg, p=0.06). Similarly, we found that the hormone simulated pregnancy treatment increases *Ttr* expression while there is a significant drop in *Ttr* following estrogen withdrawal in these mice (**Fig. 6D;** one-way ANOVA, F(2,8)=9.92, p=0.007; Tukey’s post hoc test, HSP+EB vs. OVX-Veh, p=0.008; HSP+EB vs. HSP+Veh, p=0.022).

Considering *Ttr*’s strong response to hormonal changes, we decided to further explore the potential functional relevance of *Ttr* expression in the vHIP for structural and behavioral changes across the estrous cycle^9^. We explored cell type-specific *Ttr* expression more closely and found that *Ttr* expressing cells are predominantly excitatory neurons, and their numbers increase significantly in proestrus compared to diestrus (**Extended Data Fig. 11D**). Therefore, we designed an experiment in which we overexpressed *Ttr* in excitatory neurons of the vHIP in cyclical female mice, and then tested these mice and their eGFP-expressing controls on anxiety-and depression-related behavioral tests and dendritic spine density (**Fig. 6E-F**). We tested all mice in the low-estrogenic, diestrus phase of the cycle to address the hypothesis that *Ttr* overexpression in vHIP excitatory neurons may be sufficient to shift mouse behavior and structural plasticity in a way that would be more representative of high-estrogenic hormonal state. Indeed, overexpressing *Ttr* in excitatory vHIP neurons increased time spent in the open arms of the elevated plus maze (EPM; one-tailed Mann-Whitney *U* test, p=0.002) and decreased immobility time and the number of immobility episodes in the forced swim test (FST; unpaired one-tailed *t*-test, p < 0.05) in diestrus females, to a proestrus-like state^9^, without affecting their overall activity levels (**Fig. 6E**). No significant effects were found in the open field test (data not shown). With Golgi-Cox staining, we found that *Ttr* overexpression increased dendritic spine density in low estrogenic, diestrus females (**Fig. 6F**; unpaired one-tailed *t*-test, p<0.0001). These data confirm the functional role of ovarian hormone-dependent changes in *Ttr* expression for brain structural plasticity and anxiety-and depression-related behavior.

## Discussion

Here we provide the first sex-specific, estrous cycle-dependent, and peri-partum-resolved map of the mouse brain at single cell resolution. While neuroscience has been male-centric^38,39^, stress-related disorders anxiety and depression are two times more prevalent in women and peak during reproductive transitions^40^, thus the female-focused view provided here is critical to understand sex-specific neurobiology and disease risks^41^. In particular, we find dynamic changes in the cellular composition of the vHIP, including in the neural stem cells of the dentate gyrus (DG), which are likely to mediate the increase in neurogenesis in the high-estrogenic phase of the estrous cycle^28^ and during pregnancy^42^. Adult hippocampal neurogenesis provides a substantial degree of structural and functional plasticity in the brain^27^ and it has been implicated in the antidepressant response^43^. Thus, our data provide a critical molecular insight into female-specific neuroplasticity and can inspire the development of new treatments.

Further, our study shows that the main group of cells affected by ovarian hormone shifts is, unsurprisingly, excitatory neurons that mediate changes in synaptic function and behavior. However, we are able to classify them into spatially relevant excitatory neurons including those of CA1, CA2, CA3, and DG subfields and show that each of them responds differently across reproductive transitions. CA2 cells of the vHIP were not detected in previous RNA-seq studies^16,20,44^ so we molecularly define those cells for the first time in this hippocampal subregion of mice. Furthermore, we show that while DEGs are largely limited to excitatory neurons, DARs are much more extensive including within excitatory neurons and across all cell types. We provide evidence that chromatin changes during the estrous cycle serve as a mechanism through which ovarian hormones prime the genome for future gene expression changes if the need for this arises such as during pregnancy or perhaps during stressful events. This is a likely molecular “preparatory” mechanism that allows brain cells to quickly adapt to changes in endogenous and exogenous environments, as previously shown for memory formation^45^. In this case, it allows adaptation to reproduction. However, we also show that beyond this physiological role, these chromatin changes could also prime the brain for psychopathology, particularly for depression, in individuals with genetic predisposition.

Finally, our study reveals the thyroid hormone transporter transthyretin (*Ttr*) gene as the major candidate gene responsive to both ovarian hormone shifts, across the estrous cycle and peripartum. Thyroid hormone dysregulation can affect multiple domains of brain function including a significantly increased risk for depression^46^, and both hypothyroidism and depression affect women more frequently than men^47^. A recent study showed reduced plasma protein levels of transthyretin in major depressive disorder (MDD)^48^. Other studies have shown that thyroid hormones can augment antidepressant response to selective serotonin reuptake inhibitors^49^ and a selective thyroid hormone beta receptor (TRβ) agonist is currently being tested as an adjunctive treatment for MDD^50^. None of these studies, however, took sex as a biological factor in their designs and drug development, failing to address increased female risk for depression and opportunities for more precise treatment strategies that may come with this knowledge. Our study, on the other hand, confirms the functional role of ovarian hormone-driven changes in *Ttr* expression for brain structural plasticity and anxiety-and depression-related behavior. These results open the possibility to target downstream targets of estrogen, including thyroid hormone signaling, as a treatment option for hormone-sensitive periods in women. These sex-informed strategies for treatments and prevention may revolutionize psychiatry but the comprehensive understanding of the female brain needs to be part of every discovery particularly for female-biased brain disorders.

## Data availability

Data supporting the key findings of this study are available within the article and its Supplementary Information files or from the corresponding author upon request. Correspondence and requests for materials should be addressed to M.K. and M.S.

## Acknowledgements

This work was supported by the National Institutes of Health award R01MH123523 (to M.K.). We would like to thank David Reynolds and Nickoli Parkinson for their assistance with single-cell multiome assay and next generation sequencing, respectively.

## Author contributions

M.S. and M.K. designed the study; L.D., M.T., L.O., and A.M., performed experiments. A.K.S.K., D.R., and M.S. performed analyses of the single cell data. L.D. and L.O. performed statistical analyses of gene expression, behavioral, and structural data. L.D., A.K.S.K., D.R., M.S. and M.K. interpreted the data. L.D., A.K.S.K., and D.R. constructed the figures. M.K. wrote the main article. L.D. and M.S. wrote the methods section. M.K. conceived and directed the project. All authors commented on and approved the final version of the paper.

## Materials and Methods

### Animals

For this study we used six separate cohorts of C57BL/6J mice, each dedicated to a distinct experimental paradigm. The **first cohort** was used for single-nucleus multiome (snMultiome) analysis across the estrous cycle and sex and consisted of N=12 female and N=6 male mice obtained from Jackson Laboratory at 6 weeks of age (**Fig. 1A**). After a 2-week habituation period, the estrous cycle of females was monitored across three consecutive cycles (8-11 weeks of age; see *Estrous cycle monitoring*). At 11 weeks, females were classified as proestrus (Pro; high estradiol, low progesterone) or diestrus (Die; low estradiol, high progesterone), chosen to approximate the human follicular and luteal phases, respectively, and ventral hippocampi (vHIP) were collected for snMultiome profiling (see *Single-nucleus multiome assay*). The **second cohort** was used to generate the snMultiome dataset across pregnancy and postpartum and was comprised of N=6 female and N=6 male mice, obtained from Jackson Laboratory at 6 weeks of age. After a 2-week habituation period, females were paired with males (1:1) for ten consecutive days. Pregnant females were then housed individually in standard laboratory cages to allow undisturbed delivery (see *Peripartum determination*), and vHIP tissue was collected at late gestation (gestational day GD 18) or on postpartum day 1 (within 24 h of delivery) for multiome analysis (**Fig. 5A**). The **third and fourth cohorts** were used to examine the effects of overexpressing *Ttr* in excitatory vHIP neurons during the diestrus phase (**Fig. 6E-F**). The **third cohort** consisted of N=24 female mice obtained at 6 weeks of age and following habituation, at 8 weeks of age mice underwent bilateral stereotaxic viral injections (see *Stereotaxic surgery*) into the vHIP to deliver either an AAV9 vector expressing Ttr-IRES-eGFP under the CamkII promoter (TTR group) or a matched control AAV9 expressing eGFP alone (Control group, **Fig. 6E**). After surgery, mice were allowed to recover for 3 weeks to ensure stable transgene expression while estrous cycles were monitored daily. Behavioral testing was performed between 11 and 13 weeks of age (see *Behavioral testing*), and mice were sacrificed at 14 weeks; whole brains were collected and preserved for cryosectioning (see *Immunohistochemistry*) to verify viral targeting in all animals included in behavioral analyses. The **fourth cohort** consisted of N=12 female mice that underwent the same viral manipulation (at 8 weeks of age) and was used to examine the effects of *Ttr* overexpression on dendritic spine density in vHIP neurons during diestrus (**Fig. 6F**). These animals were euthanized at 11 weeks of age and processed for Golgi-Cox staining (see *Golgi-Cox staining and dendritic spine analysis*). The **fifth and sixth cohorts** underwent ovariectomy (OVX) followed by either cyclical estradiol-replacement (**Fig. 6C**) or hormone-simulated pregnancy (HSP, **Fig. 6D**) manipulations for gene expression analyses. The **fifth cohort** comprised N=34 female mice obtained from Jackson Laboratory at 6 weeks of age. After a 2-week habituation period, mice underwent bilateral OVX or sham surgery at 8 weeks (see *Ovariectomy*). OVX mice were subsequently assigned to estradiol-replacement or vehicle conditions according to the experimental design (see *Estradiol-replacement*). The **sixth cohort** consisted of N=12 female mice obtained from Jackson Laboratory at 6 weeks of age. After a 2-week habituation period, mice underwent bilateral ovariectomy at 8 weeks old. OVX mice were subsequently assigned to hormone simulated pregnancy (HSP) or vehicle conditions according to the experimental HSP protocol (see *Hormone-simulated pregnancy*). All mice from the **sixth cohort** were euthanized on day 25 of the HSP treatment and the vHIP was dissected for gene expression analysis (see RNA isolation and *gene expression analysis*). Across all cohorts, animals were housed in same-sex cages (n=3-5 per cage) under a 12:12 h light–dark cycle with *ad libitum* access to food and water. Housing rooms were maintained at 21 °C with 30-70% humidity. Mice were euthanized by cervical dislocation followed by rapid decapitation, and whole brains or vHIP tissue were dissected immediately. All samples were stored at −80 °C until further processing. All animal procedures were approved by the Institutional Animal Care and Use Committee at Fordham University.

### Estrous cycle monitoring

Estrous cycle was assessed by daily vaginal cytology, a standard approach for identifying cycle stage^1,2^. To collect samples, 100 µl of 0.5X PBS was drawn into a transfer pipette and gently expelled at the vaginal opening to obtain exfoliated cells by lavage. The resulting suspension was placed onto glass slides and air-dried at room temperature for approximately two hours. Dried smears were stained with 0.1% crystal violet, rinsed, and examined under a light microscope. Cycle stage was determined based on the relative abundance of nucleated epithelial cells, cornified epithelial cells, and leukocytes^1^. Proestrus smears contain predominantly nucleated epithelial cells, estrus smears are characterized by cornified cells, and metestrus and diestrus contain mixtures of epithelial cells and leukocytes, with diestrus showing the highest proportion of leukocytes. Smears were collected daily across two weeks (three full cycles) to establish individual cycling patterns and to enable accurate prediction of cycle stage for molecular and behavioral experiments. Cycle assignments were verified on the day of behavioral testing or tissue collection. Animals showing irregular cycling patterns were excluded. Prior work in our laboratory confirmed that cytological staging corresponds to expected ovarian hormone profiles in serum and hippocampal tissue for proestrus (high estradiol, low progesterone) and diestrus (low estradiol, high progesterone) mice^1^.

### Peripartum determination

Pregnancy was confirmed by the presence of a vaginal plug and detection of spermatozoa in vaginal smears, together with monitoring of daily body-weight gain. Pregnant females were then randomly assigned to either the pregnancy (Preg) or postpartum (PP) group. Females in the pregnancy group were euthanized on gestational day 18 (GD18) between 9:00 and 11:00 a.m.; this time point was selected because circulating estradiol levels are known to peak in late gestation^3^. For the postpartum group, females were allowed to deliver naturally without disturbance. Dams were euthanized on postpartum day 1 (PP1), corresponding to the first day after delivery, when estradiol levels undergo a sharp physiological decline^4^. Ventral hippocampal tissue was collected immediately after euthanasia for downstream analyses.

### Single-nucleus multiome assay

Six (for the estrous cycle/sex experiment) or three (for the peripartum experiment) mice per experimental group were included in the snMultiome analysis. For the estrous cycle/sex experiment bilateral ventral hippocampi were pooled from two animals for each biological replicate (N= 3 replicates or 6 mice/group), while for the peripartum experiment a single animal for each biological replicate was used (N=3 replicates or 3 mice/group). Frozen ventral hippocampal (vHIP) tissue was processed in batches with balanced group distribution to minimize technical batch effects (three batches for the estrous-cycle cohort and two for the peripartum cohort). Nuclei were isolated using a previously established sucrose-gradient-based protocol^5^. Briefly, tissue was homogenized in ice-cold lysis buffer (0.32 M sucrose, 10mM Tris-HCl pH 8.0, 5 mM CaCl_2_, 3 mM magnesium acetate, 0.1 mM EDTA, 0.1% Triton X-100) supplemented with 1 mM DTT and RiboLock RNase inhibitor (2.5 µL/mL; ThermoFisher) using a glass Dounce homogenizer. Homogenates were layered over 1.8 M sucrose solution and nuclei were isolated by ultracentrifugation (101,814 × g, 1 h, 4 °C; Beckman ultracentrifuge). After centrifugation, the supernatant was carefully removed and nuclei pellets were resuspended in wash buffer consisting of DPBS supplemented with 10% BSA and RiboLock RNase inhibitor (25 µL/mL). Nuclei were transported on ice to the Albert Einstein College of Medicine Genomics Core for 10x Genomics processing. Upon arrival, nuclei were pelleted (500 rcf, 5 min, 4 °C), resuspended in nuclei resuspension buffer (1 mM DTT, nuclease-free water, 1x Nuclei Buffer 10x Genomics, and 25 µL/mL RiboLock RNase inhibitor), and subjected to nuclei counting and quality assessment. For each sample, 10,000 nuclei were loaded into the 10x Genomics Chromium Next GEM Single Cell Multiome ATAC + Gene Expression platform to generate parallel gene-expression (GEX) and chromatin-accessibility (ATAC) libraries. Libraries were prepared following the manufacturer’s protocol (10x Genomics) and quantified using the Qubit High-Sensitivity DNA kit (Life Technologies). Equimolar library pools were sequenced on a NovaSeq X Plus (Illumina), with GEX and ATAC libraries sequenced on separate lanes. Sequencing yielded an average depth of ~30,000 reads per nucleus for GEX and ~48,000 reads per nucleus for ATAC using paired-end 150-bp reads (PE150).

### Single-nucleus multiome data processing and quality control

#### Preprocessing

Raw FASTQ sequencing files were aligned to the mm10 mouse reference genome (2020-A release, 10x Genomics) using Cell Ranger ARC (version 2.0.2, 10x Genomics). This generated UMI-based gene expression counts and ATAC-seq fragment files for chromatin accessibility analysis. We performed downstream analyses in R (version 4.4.1) using Seurat^6–9^ (version 5.0.1) for gene expression and Signac^10^ (version 1.14.0) for chromatin accessibility data. Seurat objects were built from the matrix.h5 and fragment.tsv.gz output files with gene annotations from EnsDb.Mmusculus.v79 (doi: 10.18129/B9.bioc.EnsDb.Mmusculus.v79). We called ATAC-seq peaks using MACS2^11^ (version 2.2.7.1) and quantified reads within these peaks using Signac’s *FeatureMatrix* function. Chromosome names were converted to the UCSC format to be consistent with Cell Ranger ARC outputs. Low-quality cells with a mitochondrial gene expression percentage (<0.05), nucleosome signal < 2, and TSS enrichment score > 1 were excluded from the downstream analysis.

#### Gene expression data analysis

After quality-control filtering, we normalized each dataset using *NormalizeData* with default parameters, identified the top 3,000 highly variable genes using *FindVariableFeatures*, and scaled the expression values using *ScaleData*. To account for technical variation while preserving biological differences across conditions, we integrated samples using Seurat’s canonical correlation analysis (CCA) framework. Integration anchors were identified with *FindIntegrationAnchors* across the first 30 dimensions, and datasets were harmonized using *IntegrateData*. We performed principal component analysis with *RunPCA* and then visualized the integrated data using UMAP for downstream clustering and cell-type annotation.

#### ATAC-seq data analysis

Accessible regions detected in less than 50 cells were filtered using *FindTopFeatures*. After the filtration, normalization was performed using Term Frequency-Inverse Document Frequency (TF-IDF) transformation through *RunTFIDF* function. Dimensionality reduction was performed using singular value decomposition (SVD) via *RunSVD*. Later, the ATAC-seq data were integrated using Signac’s latent semantic indexing (LSI) framework^10^. LSI-transformed embeddings were generated independently for each sample, followed by integration through *FindIntegrationAnchors* and *IntegrateEmbeddings*.

### Multimodal integration, clustering, and annotation

The weighted nearest-neighbour (WNN) framework from Seurat, *FindMultiModalNeighbors* function, was used to integrate gene expression and chromatin accessibility data^12^. The function generated a combined neighbors graph, allocating an equal weighting of 0.5 to both the RNA-seq and ATAC-seq assays. Using this WNN graph, we performed Uniform Manifold Approximation and Projection (UMAP) for dimensionality reduction and visualization, followed by Leiden clustering for cell-type delineation with a resolution parameter of 0.8 using *FindClusters*. Initial Leiden clustering at resolution 0.8 yielded over 60 clusters, reflecting high cellular heterogeneity. To improve biological interpretability, we performed an annotation-level consolidation of selected clusters. Clusters were evaluated and manually merged when they exhibited (i) highly similar transcriptional profiles based on shared top DEGs, (ii) proximity and continuity in the WNN-based UMAP embedding. Importantly, this consolidation step did not involve re-clustering or modification of dimensionality reduction; cluster merging was performed solely during the final cell-type annotation stage. A two-step annotation method was used to identify the cell types of clusters: (a) using the Allen Brain Institute’s MapMyCells application^13^, where gene expression matrices were first converted from Seurat objects to h5ad format compatible with MapMyCells requirements (cell-by-gene matrix with species-matched gene names), then mapped against the 10X Whole Mouse Brain CCN20230722 reference atlas using hierarchical mapping algorithms (**Extended Data Table 1A**), and (b) manual curation based on top 50 differentially expressed gene markers identified through *FindAllMarkers* (Wilcoxon rank-sum test, **Extended Data Table 1B**), cross-referenced with established gene signatures from the Allen Brain Atlas^14^ and published literature^15–27^ (**Extended Data Table 1C**).

### Differentially expressed genes (DEGs) and differentially accessible regions (DARs)

The analyses of differentially expressed genes (DEGs) and differentially accessible regions (DARs) were performed using Seurat’s *FindMarkers* function. Gene expression changes were assessed with the Wilcoxon rank-sum test, while chromatin accessibility differences were evaluated using the likelihood ratio test. We used permissive initial filtering (log_2_ fold-change threshold = 0.001, minimum cell percentage = 0.05) to capture regulatory changes broadly, then applied criteria to identify biologically relevant features: adjusted p-value < 0.05 and average log_2_ fold-change >0.1 for DEGs and >0.58 for DARs (1.5-fold change). DARs were mapped to their nearest genes using Signac’s *ClosestFeature* function. We performed systematic comparisons of gene expression and chromatin accessibility across physiological states at both broad and semi-broad cell types identified. Within the diestrus-proestrus-male dataset, we performed three pairwise comparisons: (i) diestrus (vs) proestrus, (ii) diestrus (vs) male, and (iii) proestrus (vs) male. In the pregnancy-postpartum dataset, we compared pregnancy (vs) postpartum samples respectively.

### Cell proportion analysis

Cell proportion analysis to detect changes in cellular composition was performed across conditions at three different levels of resolution: (i) clusters obtained from Seurat’s clustering (resolution = 0.8), (ii) broad cell type categories, and (iii) semi-broad cell type groups. Cell type abundances were compared using the *scProportionTest*^*28*^, Bioconductor R package, with permutation-based testing (*sc_utils* and *permutation_test* functions, with 10,000 permutations). We made pairwise comparisons between diestrus (vs) proestrus, proestrus (vs) male, and diestrus (vs) male in the estrous cycle/sex dataset, and between pregnancy (vs) postpartum samples in the peripartum dataset, respectively.

### Enrichment analysis

The biological relevance of DEGs, or genes annotated to DARs, was determined using gProfiler^29^ to assess overrepresentation of genes belonging to terms and pathways in the Gene Ontology (Biological Process, Cellular Component, and Molecular Function), KEGG, and Reactome databases. Significance was considered padj < 0.05 using the g:SCS algorithm for multiple testing correction^29^ and results were plotted using the ComplexHeatmap package in R^30^. For inhibitory neurons and glia, which had fewer DEGs across group comparisons, enrichment was performed using gene set enrichment analysis (GSEA^31^, FDR < 0.10) and a publicly available gene set file (https://download.baderlab.org/EM_Genesets/current_release/Mouse/symbol/Mouse_GO_AllPathways_noPFOCRno_GO_iea_November_03_2025_symbol.gmt). For the overlap of genes annotated to Die-Pro DARs with Preg-PP DEGs, enrichment was performed using gProfiler (FDR < 0.05) with the same GMT file. Significant results from these analyses were plotted in Cytoscape^32^ using the EnrichmentMap app^33^ and clustered using AutoAnnotate^34^. Motif enrichment near DEGs was performed using gProfiler queries against the TRANSFAC database^35^ which tests for overrepresentation of transcription factor motifs within 1kb of the transcription start site of the DEG. Motif enrichment within DARs was performed using Homer^36^. To assess whether genes annotated to DARs had genetic associations with brain disorders in humans, we queried the DisGeNET^37^ and PheWeb^38^ databases using the EnrichR platform^39^.

### Adeno-associated viruses

Recombinant AAV9 vectors were used to overexpress Ttr in ventral hippocampal excitatory neurons, selected for their high transduction efficiency in brain tissue^40^. The experimental vector carried the Ttr transgene followed by an IRES-eGFP cassette, allowing coordinated expression of Ttr and eGFP from a single construct. The control vector expressed eGFP alone. Both vectors were driven by the CamkII promoter and included a woodchuck hepatitis virus post-transcriptional regulatory element (WPRE) used as an enhancer to increase the expression of the transgene. All viral preparations were obtained from Vector Biolabs (Ttr overexpression: 70700-S, eGFP control: 70600-Pre) and supplied in PBS with 5% glycerol at a titre of 10^13^ genome copies/mL.

### Stereotaxic surgery

Mice were anaesthetized with isoflurane delivered via an induction chamber and maintained under continuous isoflurane anesthesia (1.5-2%) throughout the procedure. Experimental or control AAV9 solutions (N=12 per group for behavior; N=5 per group for Golgi analyses) were injected bilaterally into the ventral hippocampus using a Nanoject III microinjector coupled to a glass pipette (15 μm tip diameter). Stereotaxic coordinates relative to bregma were: anteroposterior (A/P) −2.95 mm, mediolateral (M/L) ±2.85 mm, and dorsoventral (D/V) −3.85 mm^41^. A total volume of 200 nL was delivered per site (10 pulses x 20 nL, 15-s intervals). The dose was selected based on a dose-optimization experiment, in which 200 nL and 400 nL viral injections resulted in comparable but dose-dependent increases in *Ttr* expression (11.9-fold and 14.8-fold, respectively) relative to control (**Extended Data Fig. 12**). Post-operative care included monitoring of body weight, inspection of surgical sites, and administration of carprofen for analgesia. Mice were allowed to recover for 3 weeks to ensure stable viral transgene expression prior to behavioral testing, Golgi staining, or tissue collection. Viral targeting was verified by immunohistochemistry (see *Immunohistochemistry*), and all injected mice showed correct targeting.

### Immunohistochemistry

Freshly dissected brains were rinsed in ice-cold 0.1 M PBS and fixed in 4% paraformaldehyde (PFA) in 0.1 M PBS at 4 °C for 24 h. After fixation, brains were washed in cold PBS and cryoprotected by immersion in 15% sucrose followed by 30% sucrose (both in 0.1 M PBS) for 24 h and 48 h, respectively, at 4 °C. Cryoprotected brains were then embedded, frozen in dry ice-cooled hexane, and stored at −80 °C until sectioning. Coronal sections (25 μm) containing the ventral hippocampus were cut on a rotary cryostat (Leica CM1950; Leica Biosystems) and mounted onto SuperFrost Ultra Plus slides (Fisher Scientific), then stored at −80 °C until immunostaining. For staining, slides were rehydrated in 0.1 M PBS for 30 min at room temperature, followed by blocking in 5% BSA with 0.4% Triton X-100 in PBS for 1 h. Sections were then washed in PBS-T (1% BSA, 0.4% Triton X-100 in PBS) and incubated overnight at 4 °C with rabbit anti-Ttr primary antibody (Invitrogen MA5-47192, 1:500 in PBS-T). After primary incubation, sections were washed and incubated with donkey anti-rabbit IgG secondary antibody (Invitrogen A-21207, 1:250 in PBS-T) for 2 h at room temperature in the dark. Slides were then washed again, counterstained with DAPI (1:1000 in PBS, 5 min), rinsed, and coverslipped using Mowiol 4-88 mounting medium. Negative controls lacking either primary or secondary antibody were included in each staining session to confirm that fluorescence reflected specific detection of Ttr rather than tissue autofluorescence or nonspecific secondary antibody binding.

### Behavioral testing

Mice were assigned to two groups (Ttr overexpression and eGFP Control; N=12 per group) and underwent behavioral testing between 11 and 13 weeks of age. All assessments were conducted between 9:00 and 11:00 a.m. and exclusively during early diestrus, corresponding to low estradiol and high progesterone levels. The estrous cycle was monitored daily for three consecutive cycles so that early diestrus on the day of test could be predicted. Early diestrus was confirmed by vaginal cytology on the morning of each test day. Behavioral tests were performed in the following order (from least to most stressful) with at least 24 h between sessions: open field, elevated plus maze, and forced swim test. All behaviors were video-recorded and automatically tracked using ANY-maze software (Stoelting Co., IL). Experimenters were blinded to group assignment for all scoring and analyses. For *Open-field test*, mice were placed individually in a square acrylic arena (40 × 40 × 35 cm; Stoelting) and allowed to explore the arena freely for 10 min. The arena was divided into a center zone (20 x 20 cm) and periphery. Time spent in the center and total distance traveled were quantified. For *Elevated plus maze*, the apparatus consisted of a plus-shaped maze (Stoelting) with two open arms (35 x 5 cm) and two closed arms of equal size enclosed by 15-cm walls, elevated 50 cm above the floor. Each mouse was placed on the central platform (5 x 5 cm) facing an open arm and allowed to explore for 5 min. Time spent in open versus closed arms and total distance traveled were recorded. For *Forced swim test*, mice were placed individually in 2-L glass beakers filled with 1 L of clean, room-temperature water and recorded for 6 min. Time spent immobile and number of immobility episodes were quantified using ANY-maze.

### Golgi-Cox staining and dendritic spine analysis

Whole brains of mice injected with either eGFP control or Ttr AAV (N=5/group) were rinsed with distilled water and processed using the Golgi-Cox OptimStain Kit (HitobiotecInc., #HTKNS1125) according to the manufacturer’s instructions. Briefly, brains were submerged in Golgi-Cox impregnation solution and stored in the dark at room temperature for 24 h, after which the solution was replaced and brains remained in impregnation solution for 2 weeks. Samples were then transferred to tissue-protectant solution and stored in the dark at 4 °C for 12 h; the solution was then replaced and brains remained in protectant solution for 72 h. Following impregnation, brains were frozen in dry ice-cooled hexane and stored at −80 °C until sectioning. OCT-embedded brains were cut into 100 μm coronal sections using a cryostat (Leica CM1950) and collected on gelatin-coated slides. After drying overnight, slides were stained according to the kit protocol and mounted with DPX (Sigma-Aldrich). Brightfield Z-stack images of dendrites from vHIP CA1 pyramidal neurons were acquired at 60X magnification using the Zeiss Axio Observer Microscope with Axiocam MRc Zeiss camera and Axiovision 4.8 Software (Zeiss). Spine density and morphology were analyzed using ImageJ (https://imagej.nih.gov/ij/index.html).

### Ovariectomy

Mice from Cohorts 5 and 6 underwent ovariectomy. Mice arrived at 6 weeks of age and were habituated for 2 weeks under standard housing conditions. At 8 weeks, bilateral ovariectomy or sham surgery was performed under isoflurane anesthesia. Ovaries were excised at the tip of the uterine horn via a bilateral incision; sham animals underwent the same procedure without ovary removal. Post-operative care included body-weight monitoring, wound inspection, carprofen analgesia, and Nutra-Gel (Bioserv, S4798) supplementation. The efficiency of the ovariectomy surgery was confirmed by absence of estrous cycling using vaginal cytology and by post-mortem uterine atrophy.

### Estradiol-replacement

For estradiol replacement, ovariectomized (OVX) mice (N=34 total) began receiving estradiol benzoate (EB) one week after surgery (9 weeks of age). EB was administered subcutaneously every four days for six weeks (0.2 μg or 1 μg per injection) to mimic the physiological estradiol rhythm^42^. Intact and sham-operated females were tracked by vaginal cytology and classified as diestrus or proestrus at sacrifice. All animals were euthanized at 15 weeks of age, and the ventral hippocampus was collected for gene-expression analysis. For analysis conducted in Cohort 5, diestrus and proestrus mice from sham-operated and control groups were combined, as no relevant effects of sham surgery were detected^42^.

### Hormone-simulated pregnancy (HSP)

A separate OVX cohort (N=12) was used for the HSP protocol^43^. From days 1-16, mice received daily s.c. injections of EB (0.5 μg) and progesterone (P4; 80 μg) in corn oil to simulate early pregnancy. From days 17-23, P4 was discontinued and EB was increased to 10 μg/day to simulate late pregnancy. On day 24, mice either continued EB (HSP+EB) to model late pregnancy or were switched to vehicle (HSP+Veh) to model postpartum estrogen withdrawal. Vehicle controls received corn oil throughout. All mice were euthanized on day 25 between 9:00 and 11:00 a.m., and the ventral hippocampus was dissected for gene expression analysis.

### RNA isolation and bulk qRT-PCR gene expression analysis

For estradiol replacement and hormone-simulated pregnancy models, Ttr expression was determined using bulk tissue qRT-PCR analysis. RNA was isolated from frozen ventral hippocampal samples using the AllPrep DNA/RNA Mini Kit (QIAGEN, Cat. #80204). cDNA was synthesized from 500 ng RNA (SuperScript III, Invitrogen). qPCR was performed with SYBR Green Master Mix (Applied Biosystems, #4385612) on a QuantStudio3 Real-Time PCR system. Relative expression was calculated using the 2^-ΔΔCt^ method with Ppia as the reference gene. Primer sequences were: Ttr (forward 5’-TCGCGGATGTGGTTTTCACAG-3’; reverse 5’-CTCTCAATTCTG-GGGGTTGCT-3’) and Ppia (forward 5’-GAGCTGTTTGCAGACAAAGTTC-3’; reverse 5’-CCCTGGCACATGAATCCTGG-3’).

### Statistics

Statistical analyses were performed using GraphPad Prism and R. Normality of data distribution was assessed using Shapiro-Wilk test. Candidate gene expression data (**Fig. 6C–D**) involving comparisons across multiple groups, were analyzed using one-way analysis of variance (ANOVA), followed by Tukey’s post hoc test to correct for multiple comparisons. Behavioral measures and dendritic spine density in the Ttr overexpression experiment (**Fig. 6E–F**), involving comparisons between two independent groups, were analyzed using unpaired one-tailed *t*-tests when data were normally distributed, or Mann-Whitney tests when normality was not met. One-tailed *t*-tests were used for *Ttr* overexpression experiments where a directional hypothesis was defined *a priori*. For all analyses, statistical significance was set at *p* < 0.05.

---

**Extended Data Figure 1.**
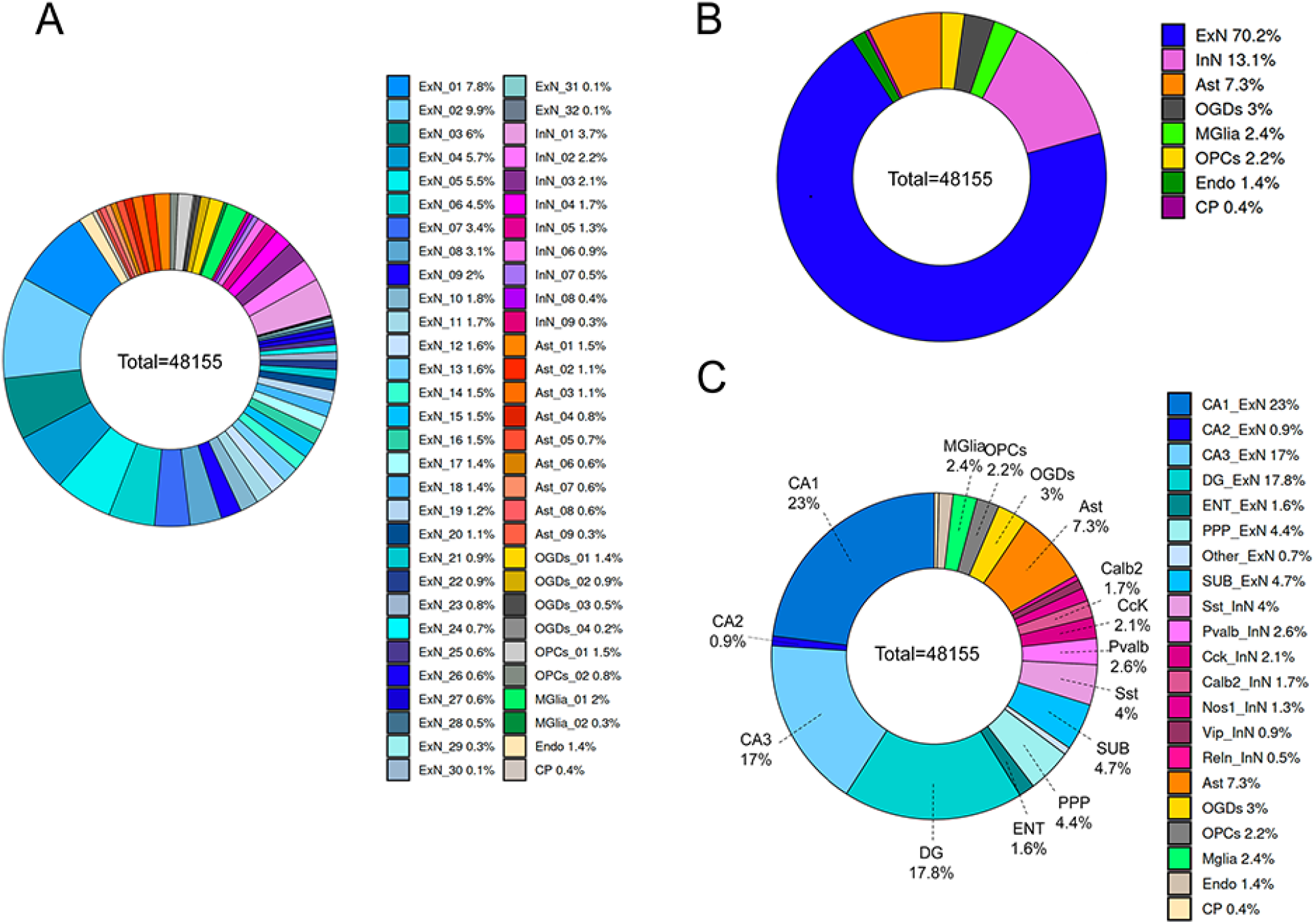
Cellular proportions in the ventral hippocampus across three levels of analysis. **A**. 60 cell clusters. **B**. Broad cell types. **C**. Semi-broad cell types. For this analysis, all nuclei across groups (N=48,155) were included. ExN, excitatory neurons; InN, inhibitory neurons; Ast, astrocytes. OGD, oligodendrocytes; MGlia, microglia; OPC, oligodendrocyte precursor cells; Endo, endothelial cells; CP, choroid plexus. DG, dendate gyrus; ENT, entorhinal cortex; PPP, prosubiculum/presubiculum/parasubiculum; SUB, subiculum, Pvalb, parvalbumin; Sst, somatostatin; Vip, vasoactive intestinal peptide; Cck, cholecystokinin; Calb2, calbindin 2; Nos1, neuronal nitric oxide synthase; Reln, reelin.

**Extended Data Figure 2.**
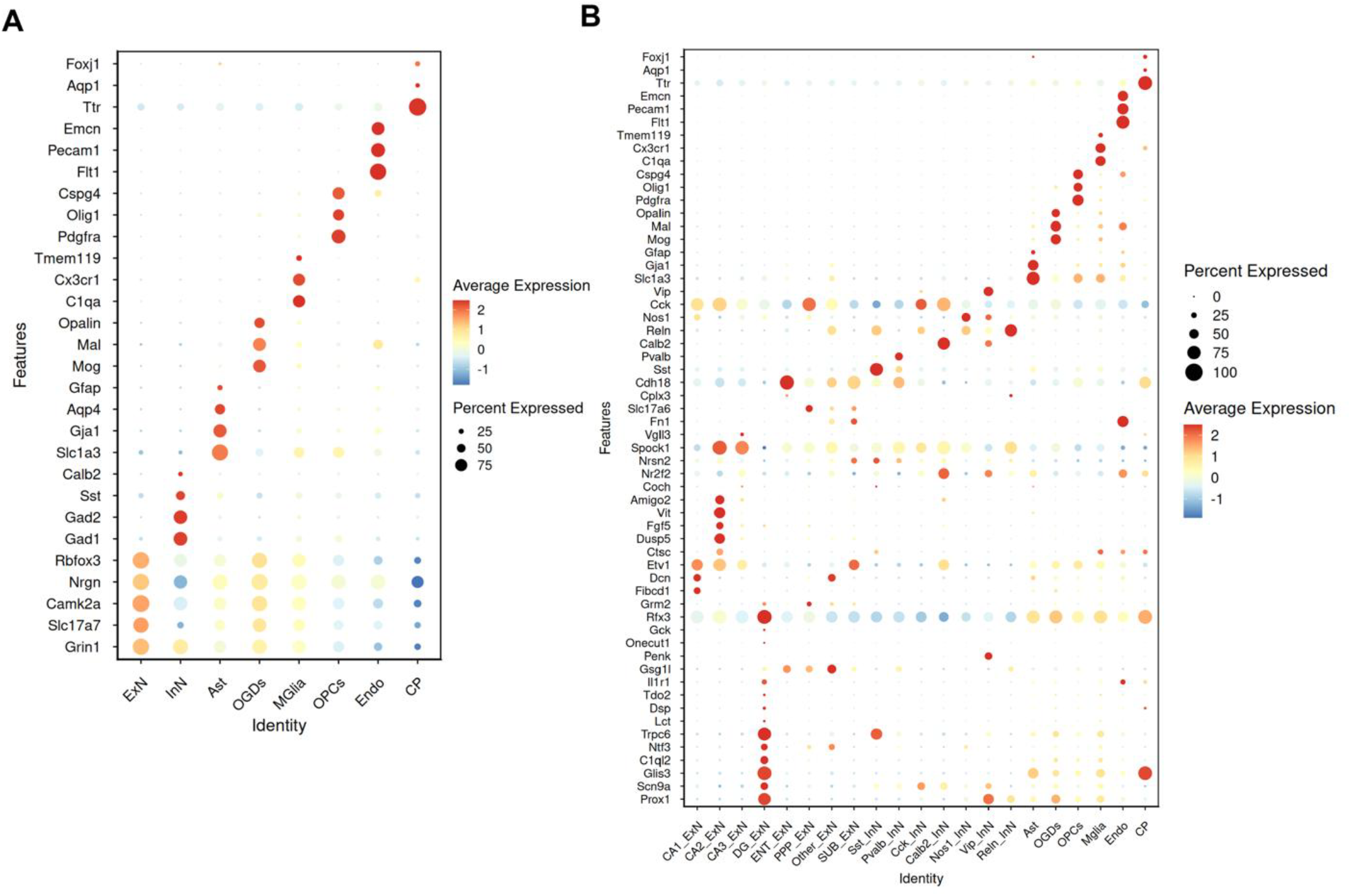
Broad and semi-broad classification of the ventral hippocampus cells. Shown are marker genes (Y axis) that were used to classify neurons into: **A**. Broad cell types of excitatory neurons (ExN), inhibitory neurons (InN), astrocytes (Ast), oligodendrocytes (OGDs), microglia (MGlia), oligodendrocyte precursor cells (OPCs), endothelial cells (Endo), and choroid plexus (CP) cells (X-axis); or **B**. subgroups of excitatory (CA1, CA2, CA3, dentate gyrus - DG) and inhibitory (Sst, Pvalb, Cck, Calb2, Nos1, Vip, Reln) neurons, along astrocytes, oligodendrocytes, microglia, oligodendrocyte precursor cells, endothelial cells, choroid plexus cells, as well as a small populations of entorhinal cortex (ENT), prosubiculum/presubiculum/parasubiculum (PPP), and subiculum (SUB) excitatory neurons (X-axis). Pvalb, parvalbumin; Sst, somatostatin; Vip, vasoactive intestinal peptide; Cck, cholecystokinin; Calb2, calbindin 2; Nos1, neuronal nitric oxide synthase; Reln, reelin. Dot size indicates the percentage of cells expressing each gene, and dot color represents average gene expression.

**Extended Data Figure 3.**
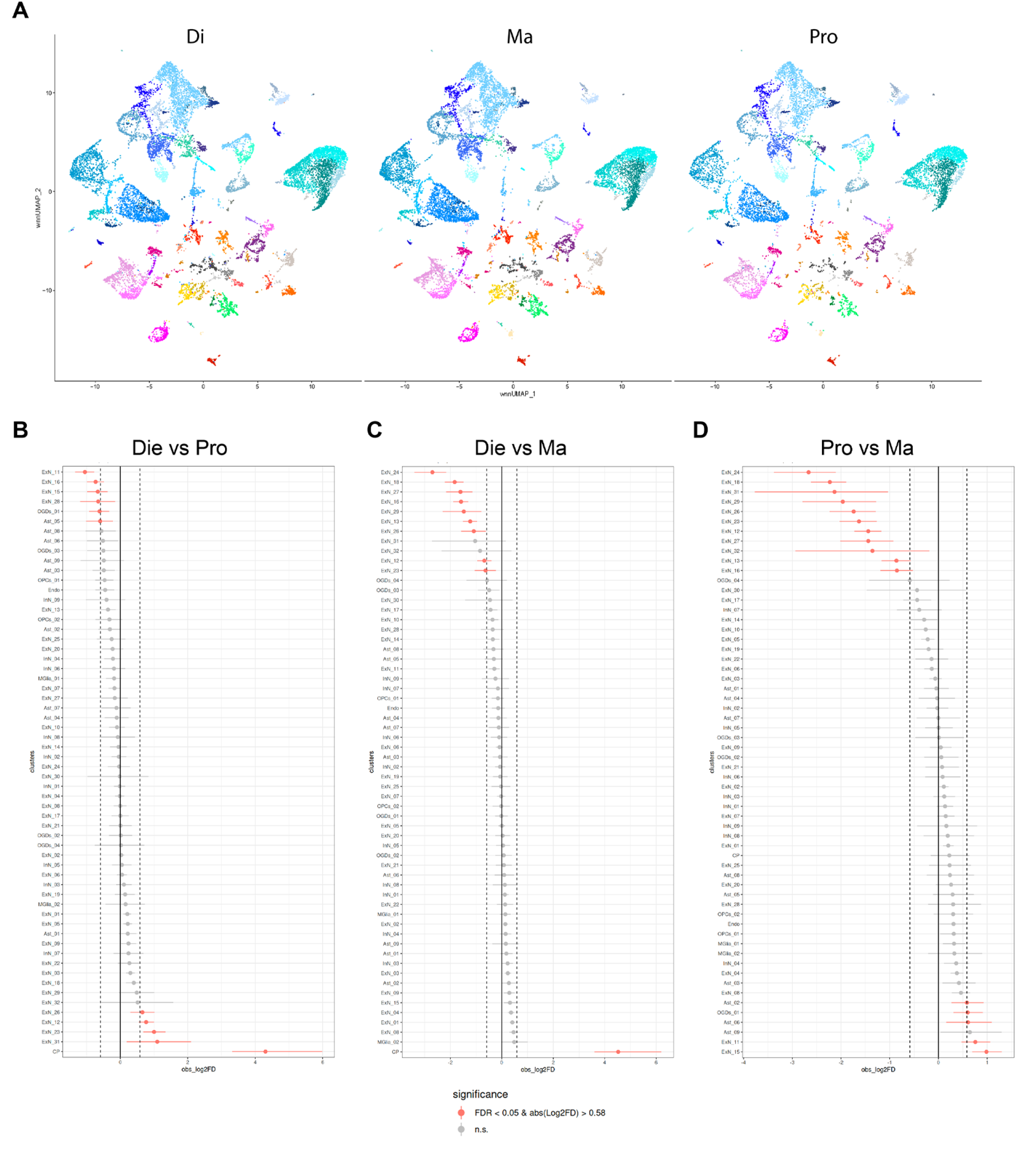
Analysis of cellular proportions in the ventral hippocampus (vHIP) across the estrous cycle and sex. **A**. UMAP shows cellular profiles of the vHIP in diestrus females (Di), proestrus females (Pro), and males (Ma). **B**-**D**. We found cellular proportion changes in all comparisons: Diestrus-Proestrus, Diestrus-Males, and Proestrus-Males, with the most significant differences found in the Proestrus-Male comparison, likely driven by estrogen rise in the proestrus stage, since we also see a significant, dynamic effect of the estrous cycle stage on vHIP cellular proportions. FDR < 0.05 was considered significant and is shown in red color. Excitatory neurons (ExN); inhibitory neurons (InN); astrocytes (Ast); oligodendrocytes (OGDs); microglia (MGlia); oligodendrocyte precursor cells (OPCs); endothelial cells (Endo); choroid plexus (CP) cells.

**Extended Data Figure 4.**
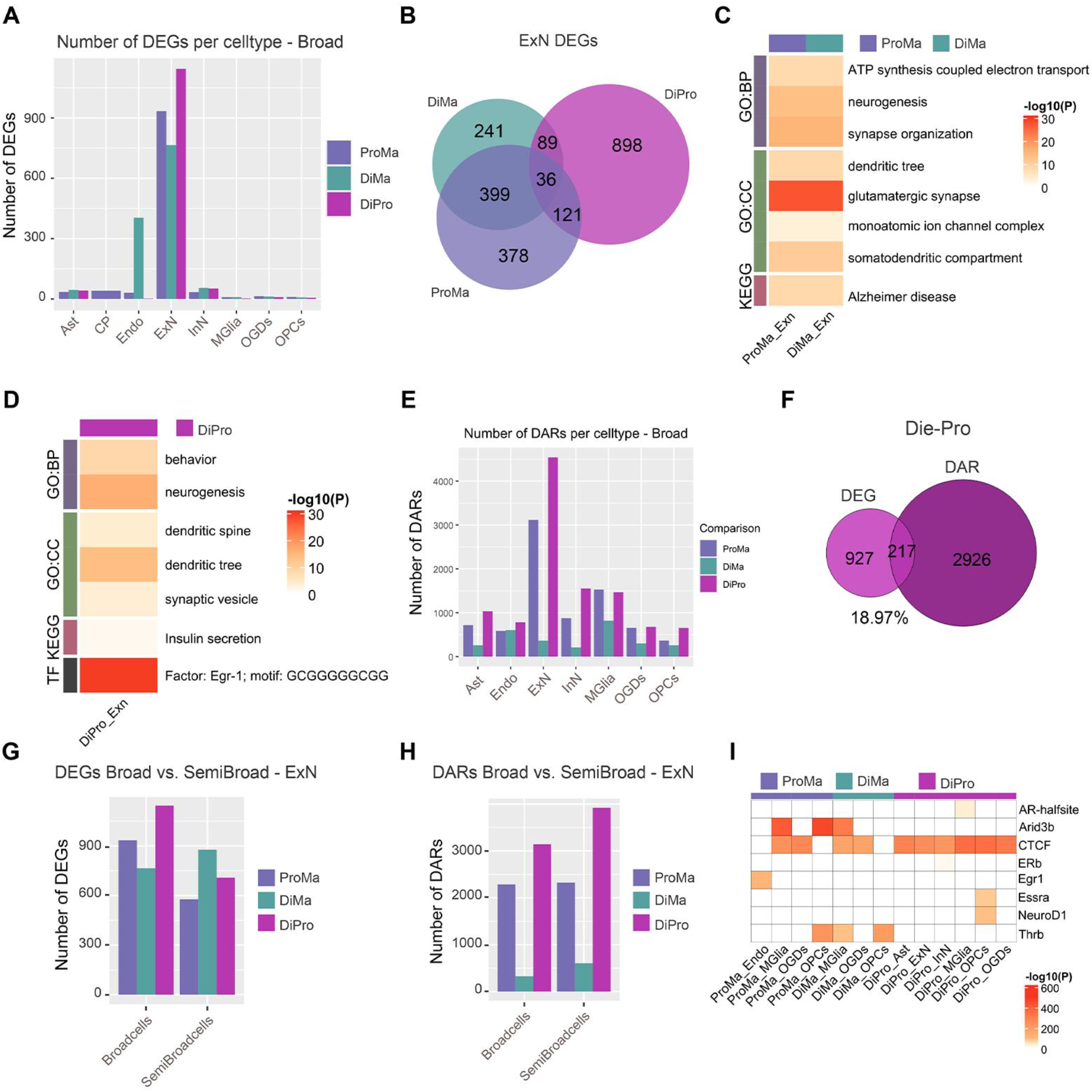
Excitatory neurons are the major cell type affected by the estrous cycle and sex. **A**. Bar graphs showing the number of differentially expressed genes (DEGs; p_adj_ < 0.05 & |log_2_fold-change| > 0.1) across broad cell types Proestrus-Male (ProMa), Diestrus-Male (DiMa), and Diestrus-Proestrus (DiPro) group comparisons. **B**. A Venn diagram showing the overlaps across the three comparisons of DEGs in excitatory neurons (ExN). Heatmaps of enrichment analysis of DEGs in broad excitatory neurons in: **C**. ProMa and DiMa comparisons (N=399, see **B**) and **D**. DiPro comparison (N=898, see **B**). **E**. Bar graphs showing the number of differentially accessible regions (DARs, p_adj_ < 0.05 & |log_2_fold-change| > 0.58) across broad cell types within ProMa, DiMa, and DiPro group comparisons. **F**. A Venn diagram showing the overlap between DEGs and genes annotated to DARs in excitatory neurons in the ventral hippocampus. Bar graphs showing the comparison of the number of DEGs (**G**) and DARs (**H**) across broad and semi-broad cell types in ProMa, DiMa, and DiPro group comparisons. **I**. A heatmap depicting the transcription factor motifs highlighted in semi-broad cell types in **Figure 3D** that are also enriched in DARs across broad cell types within any group comparison. Enrichment heatmaps (**C, D**) show statistically significant terms and pathways (p_adj_ < 0.05) from Gene Ontology (GO) Biological Process (BP), Cellular Component (CC) databases, as well as KEGG pathways and TRANSFAC motifs. Heatmap cells are colored by −log10(P_adj_) values. Excitatory neurons (ExN); inhibitory neurons (InN); astrocytes (Ast); oligodendrocytes (OGDs); microglia (MGlia); oligodendrocyte precursor cells (OPCs); endothelial cells (Endo); choroid plexus (CP) cells.

**Extended Data Figure 5.**
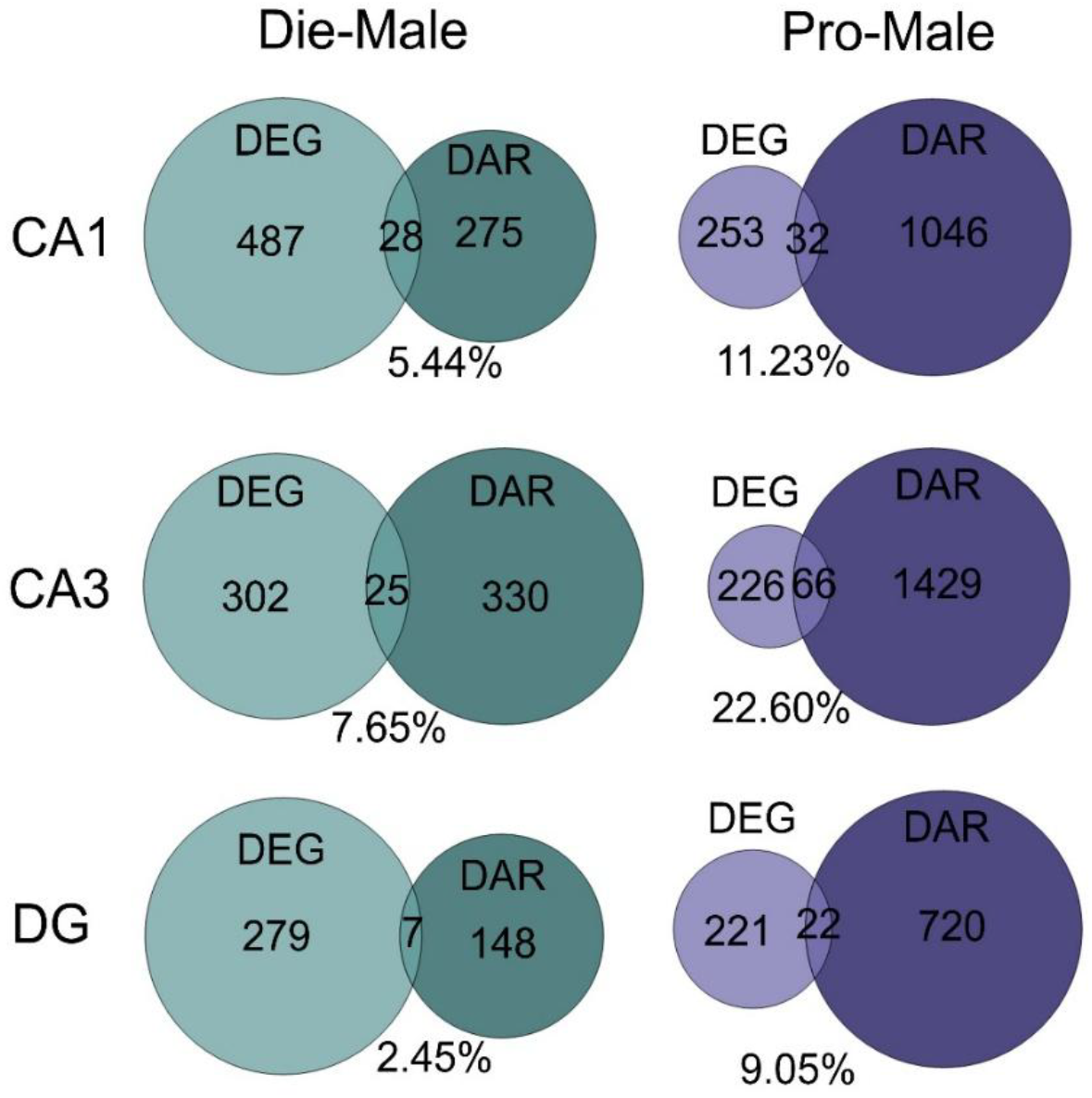
Sex-dependent overlaps between differentially expressed genes (DEGs) and differentially accessible regions (DARs) across excitatory neuron cell types. Venn diagrams show the overlap between DEGs and genes annotated to DARs in Diestrus-Male (Die-Male, left) and Proestrus-Male (Pro-Male, right) comparisons across major excitatory neurons CA1 (top), CA3 (middle), and DG (dentate gyrus; bottom).

**Extended Data Figure 6.**
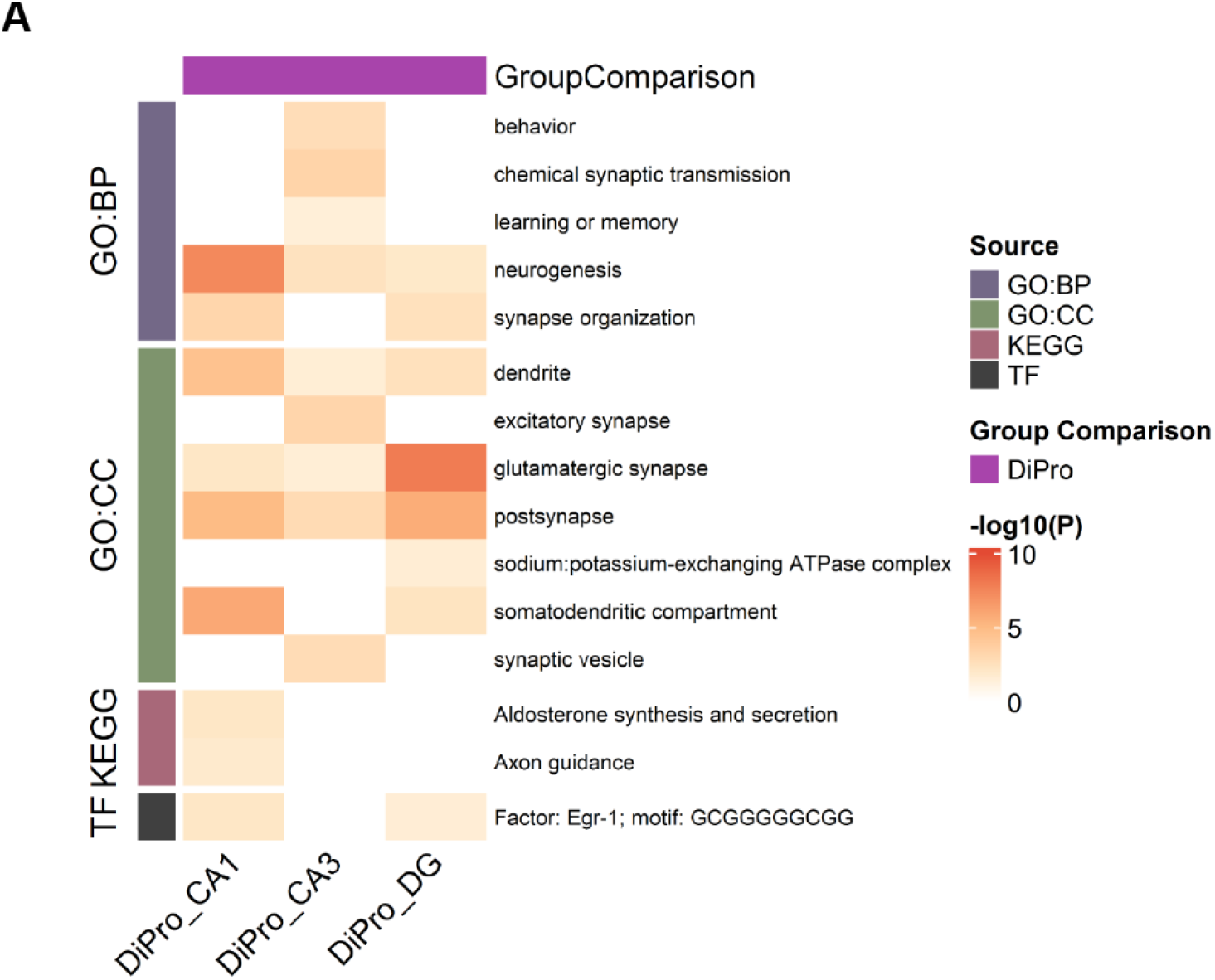
Enrichment analysis of the overlap genes between DEGs and DARs across the estrous cycle. The enrichment heatmap shows statistically significant terms and pathways (p_adj_ < 0.05) from Gene Ontology (GO) Biological Process (BP) and Cellular Component (CC) databases, as well as KEGG pathways and TRANSFAC motifs in the Diestrus-Proestrus (DiPro) comparison across ventral hippocampal CA1, CA3, and dentate gyrus (DG) cell types. Heatmap cells are colored by −log10(P_adj_) values.

**Extended Data Figure 7.**
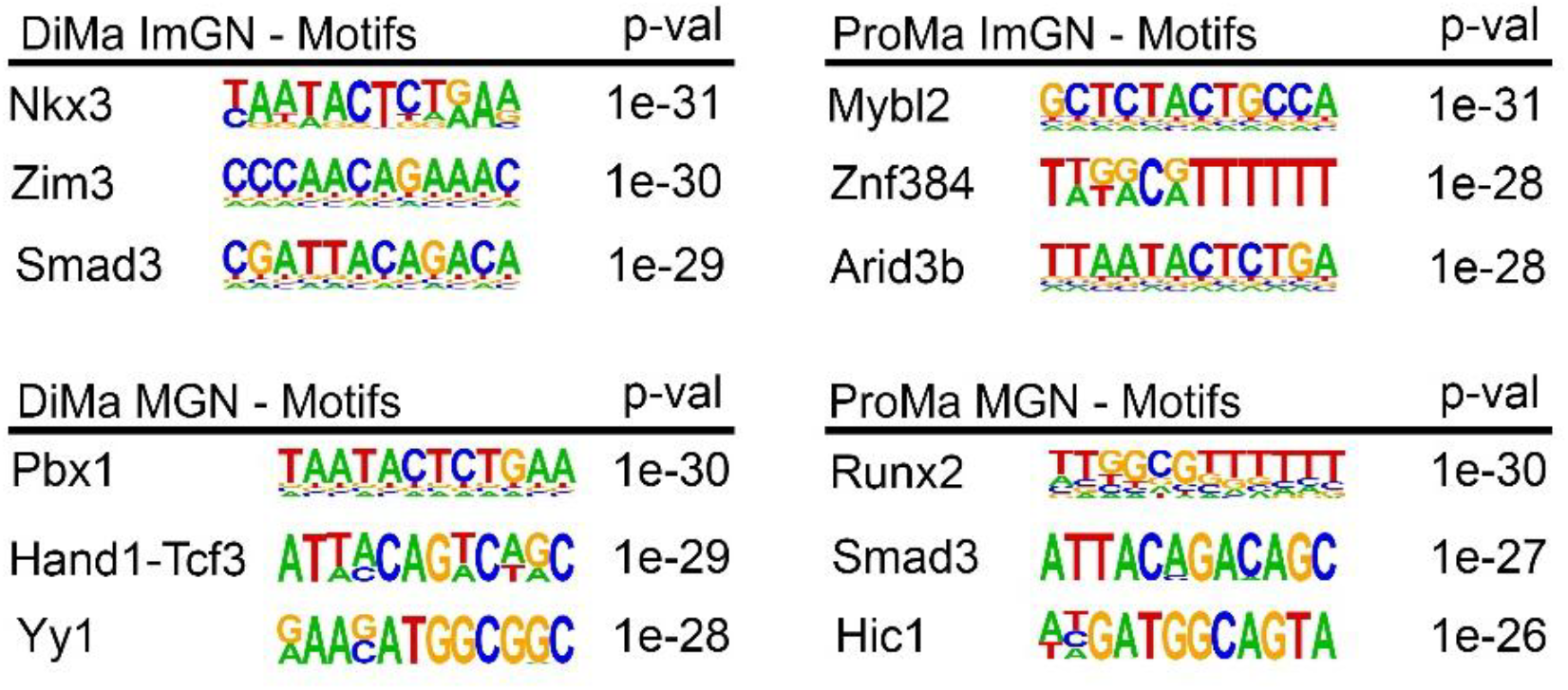
Motif analysis of chromatin changes in immature granule neurons (ImGN) and mature granule neurons (MGN) across sex. Shown are top three motifs found in the Homer analysis of single cell chromatin accessibility data in ImGN and MGN neurons in the Diestrus-Male (DiMa; on the left) and Proestrus-Male (ProMa; right) comparison.

**Extended Data Figure 8.**
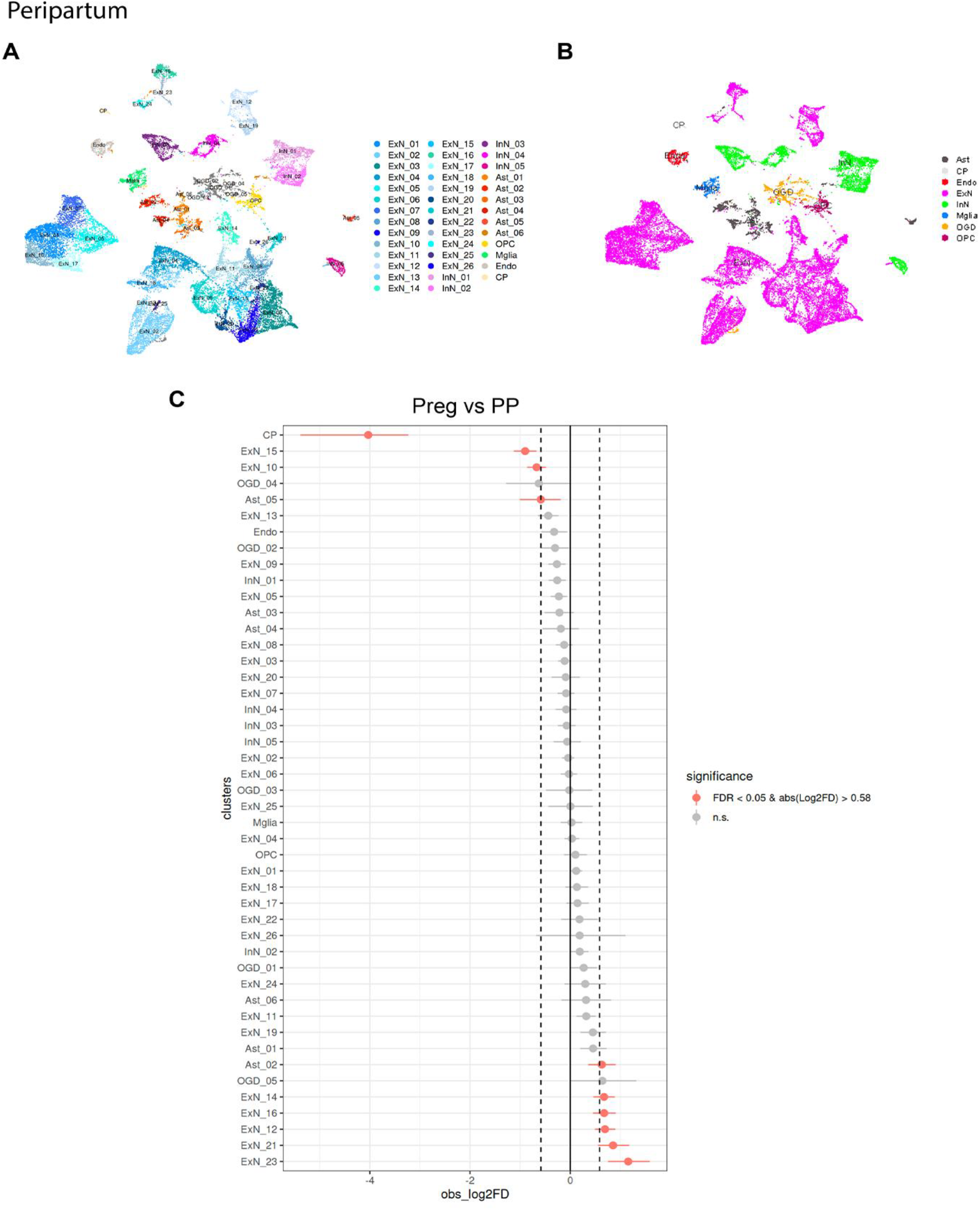
Cellular composition of the ventral hippocampus (vHIP) during the peripartum transition. Shown are UMAPs of the high-resolution 46-cluster (**A**) and broad (**B**) cell classifications of the vHIP from the single-cell multiome data derived from the female vHIP in late pregnancy (Preg; gestational day - GD 18) and postpartum (PP; N=3 biological replicates or 3 animals/group). **C**. 10 cellular clusters show changes in cellular proportions across this transition (Preg vs PP). FDR < 0.05 was considered significant and is shown in red color. Excitatory neurons (ExN); inhibitory neurons (InN); astrocytes (Ast); oligodendrocytes (OGDs); microglia (MGlia); oligodendrocyte precursor cells (OPCs); endothelial cells (Endo); choroid plexus (CP) cells.

**Extended Data Figure 9.**
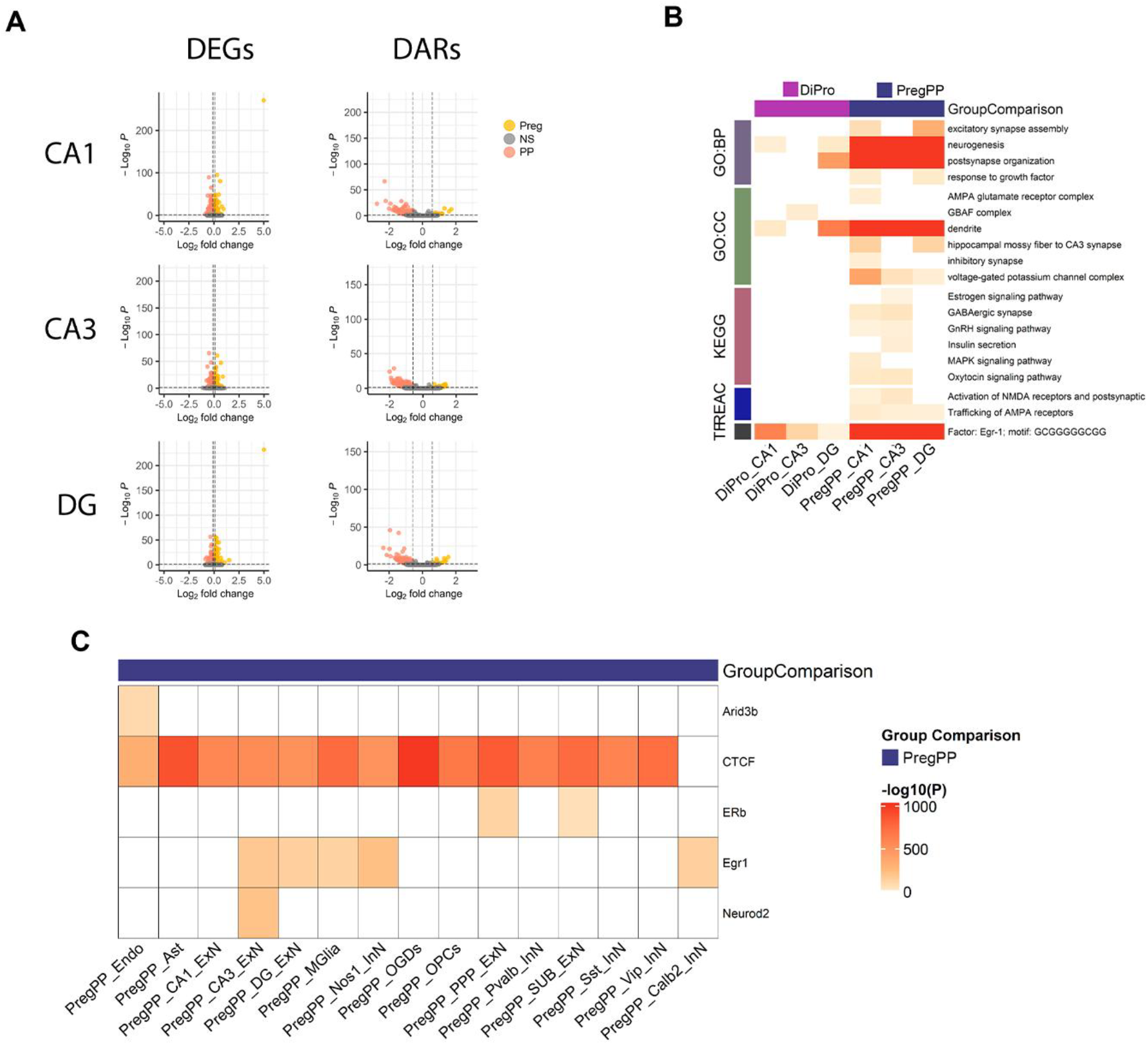
Gene expression and chromatin changes across the peripartum transition. **A.**Shown are differentially expressed genes (DEGs) and differentially accessible regions (DARs), as well as **B**. Comparison of the enrichment terms in the Diestrus-Proestrus (DiPro) and Pregnancy-Postpartum (PregPP) period, across major excitatory neuron subtypes CA1, CA3, and DG. The enrichment heatmap shows statistically significant terms and pathways (p_adj_ < 0.05) from Gene Ontology (GO) Biological Process (BP) and Cellular Component (CC) databases, as well as KEGG pathways and TRANSFAC. **C**. A heatmap depicting the transcription factor motifs highlighted in **Figure 3D** that are enriched in DARs in the pregnancy-postpartum (PregPP) comparison across all major semi-broad cell types. Heatmap cells are colored by −log10(P_adj_) values. ExN, excitatory neurons; InN, inhibitory neurons; Ast, astrocytes; OGD, oligodendrocytes; MGlia, microglia; OPC, oligodendrocyte precursor cells; Endo, endothelial cells; DG, dentate gyrus; PPP, prosubiculum/presubiculum/parasubiculum; SUB, subiculum; Pvalb, parvalbumin; Sst, somatostatin; Vip, vasoactive intestinal peptide; Calb2, calbindin 2; Nos1, neuronal nitric oxide synthase.

**Extended Data Figure 10.**
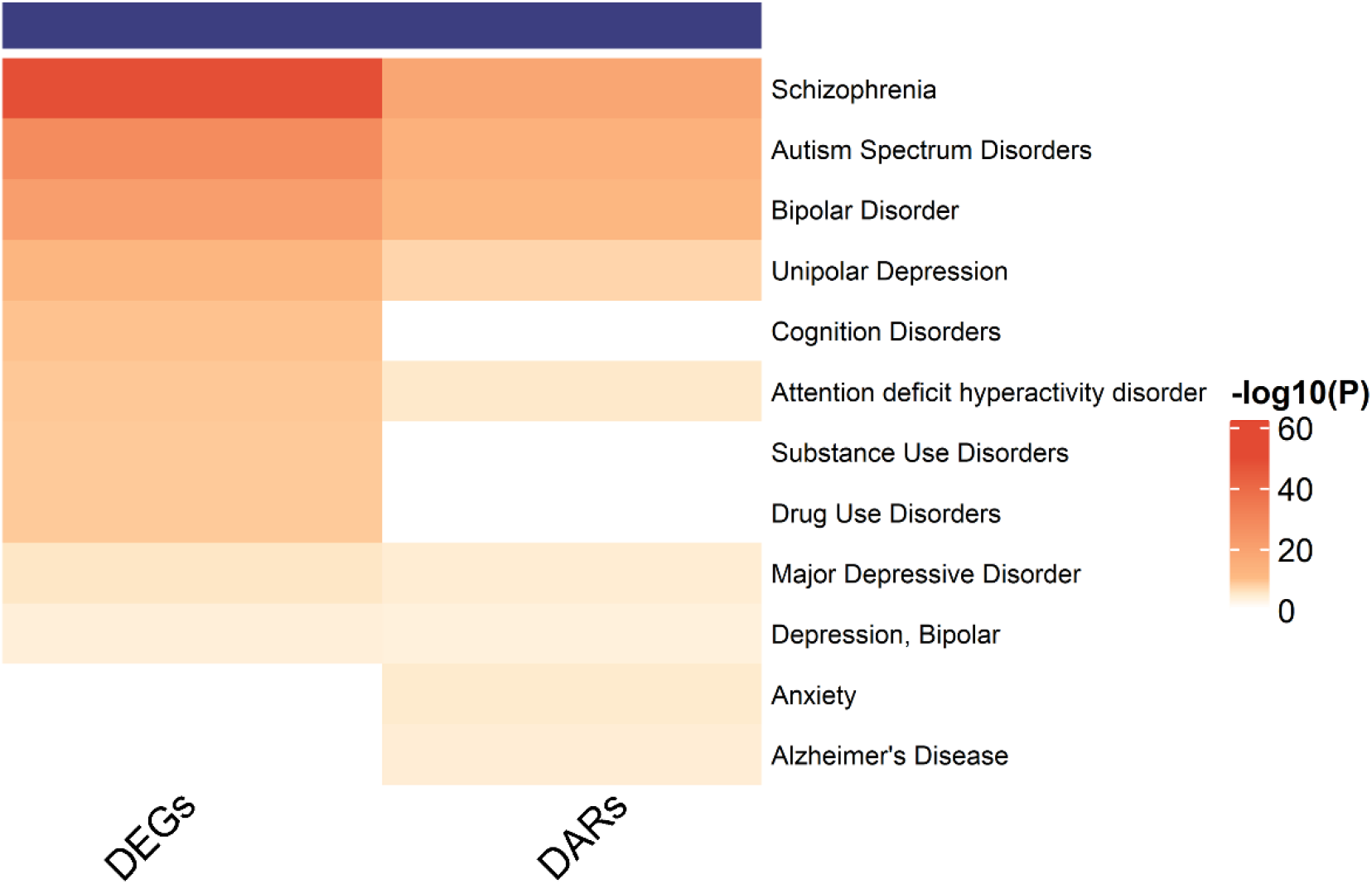
Enrichment of differentially expressed genes (DEGs) and genes annotated to differentially accessible regions (DARs) across the peripartum period for genetic associations with human brain disorders. A heatmap shows the enrichment of DEGs and genes annotated to DARs for genetic associations with human brain disorder using the DisGeNET database in the pregnancy-postpartum comparison. Heatmap cells are colored by −log10(P_adj_) values.

**Extended Data Figure 11.**
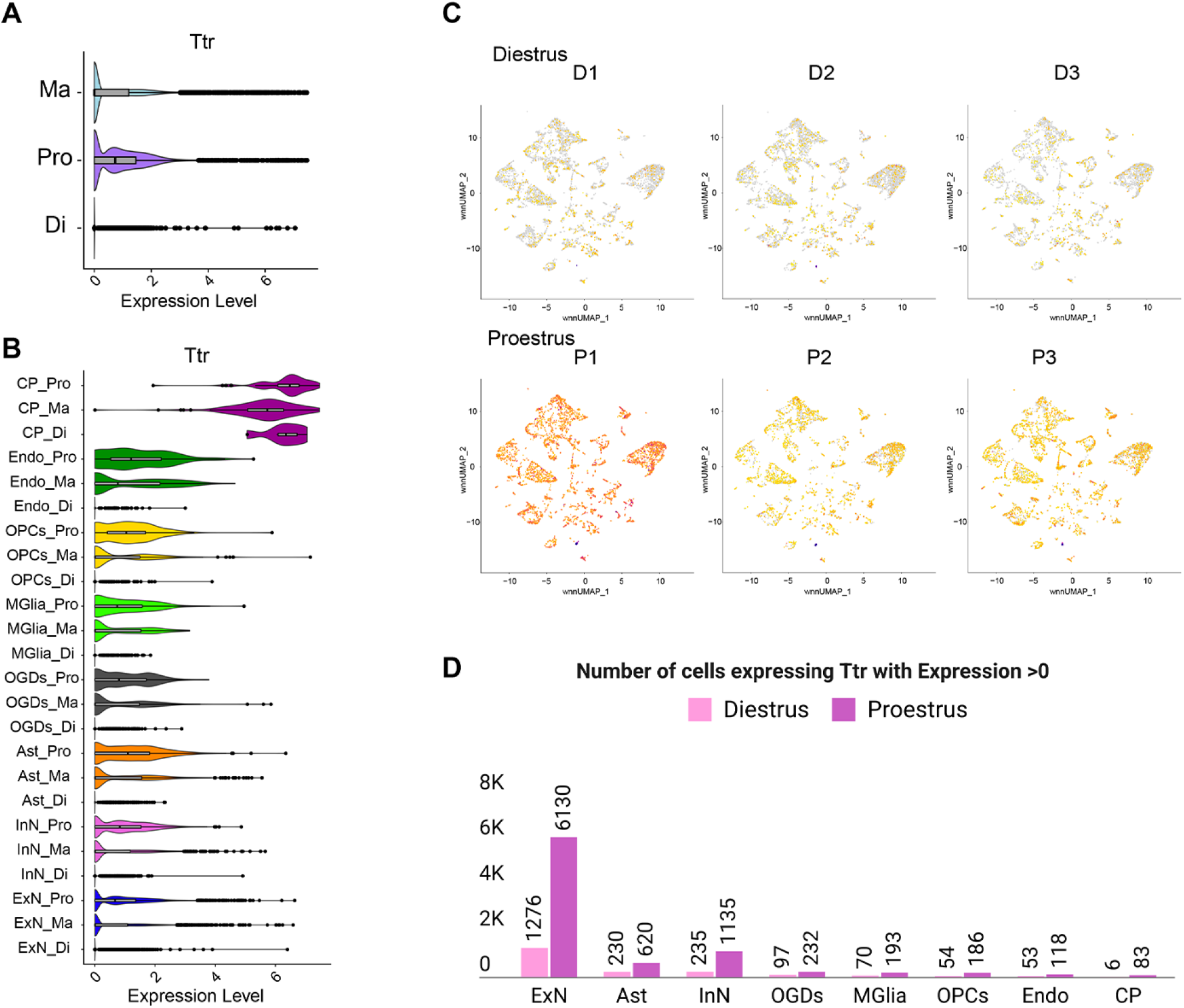
The transthyretin gene expression across the estrous cycle and sex. Higher transthyretin (*Ttr)* gene expression during proestrus compared to the diestrus stage of the cycle is clearly observed: in bulk cell comparison (**A**); across broad cell types with the exception of the CP cluster (**B**); and across replicates (**C**). Excitatory neurons are the cell type with the highest number of Ttr expressing cells, and this number increases in proestrus compared to diestrus (**D**). ExN, excitatory neurons; InN, inhibitory neurons; Ast, astrocytes. OGD, oligodendrocytes; MGlia, microglia; OPC, oligodendrocyte precursor cells; Endo, endothelial cells; CP, choroid plexus.

**Extended Data Figure 12.**
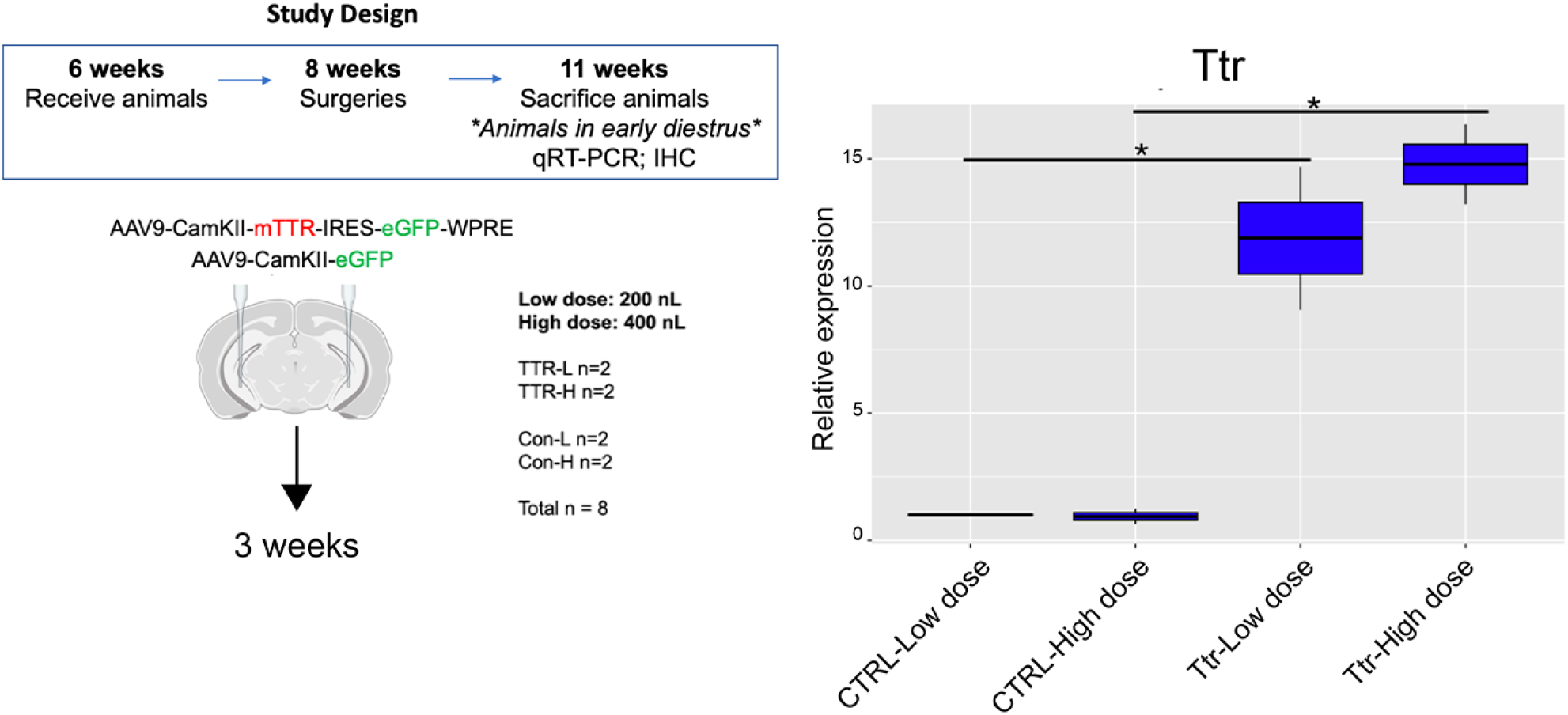
Dose-dependent validation of Ttr overexpression in ventral hippocampal excitatory neurons. Experimental design and timeline (on the left). Female mice received bilateral injections of AAV9-CamKII-mTTR-IRES-eGFP or control AAV9-CamKII-eGFP into the ventral hippocampus at 8 weeks of age and were sacrificed 3 weeks later during early diestrus. Low (200 nL) and high (400 nL) dose viral injections were used for both Ttr and control (CTRL) vectors (CTRL-Low dose, N = 2; CTRL-High dose, N = 2; Ttr-Low dose, N = 2; Ttr-High dose, N = 2; total N = 8). Tissue was collected for qRT-PCR and immunohistochemistry. On the right: qRT-PCR analysis showing dose-dependent increases in *Ttr* expression in ventral hippocampal tissue following viral overexpression. Data are presented as relative gene expression using the 2^-ΔΔCT^ method. One-way ANOVA followed by Tukey’s post hoc test; P < 0.05.

